# Heterogeneity of the group B streptococcal type VII secretion system and influence on colonization of the female genital tract

**DOI:** 10.1101/2023.01.25.525443

**Authors:** Brady L. Spencer, Alyx M. Job, Clare M. Robertson, Zainab A. Hameed, Camille Serchejian, Caitlin S. Wiafe-Kwakye, Jéssica C. Mendonça, Morgan A. Apolonio, Prescilla E. Nagao, Melody N. Neely, Natalia Korotkova, Konstantin V. Korotkov, Kathryn A. Patras, Kelly S. Doran

## Abstract

Type VIIb secretion systems (T7SSb) in Gram-positive bacteria facilitate physiology, interbacterial competition, and/or virulence via EssC ATPase-driven secretion of small ɑ-helical proteins and toxins. Recently, we characterized T7SSb in group B *Streptococcus* (GBS), a leading cause of infection in newborns and immunocompromised adults. GBS T7SS comprises four subtypes based on variation in the C-terminus of EssC and the repertoire of downstream effectors; however, the intra-species diversity of GBS T7SS and impact on GBS-host interactions remains unknown. Bioinformatic analysis indicates that GBS T7SS loci encode subtype-specific putative effectors, which have low inter-species and inter-subtype homology but contain similar domains/motifs and therefore may serve similar functions. We further identify orphaned GBS WXG100 proteins. Functionally, we show that GBS T7SS subtype I and III strains secrete EsxA *in vitro* and that in subtype I strain CJB111, *esxA1* appears to be differentially transcribed from the T7SS operon. Further, we observe subtype-specific effects of GBS T7SS on host colonization, as subtype I but not subtype III T7SS promotes GBS vaginal persistence. Finally, we observe that T7SS subtypes I and II are the predominant subtypes in clinical GBS isolates. This study highlights the potential impact of T7SS heterogeneity on host-GBS interactions.

## 1 INTRODUCTION

Type VIIb secretion systems (T7SSb) in Gram-positive organisms contribute to interbacterial competition as well as virulence by damaging host cells and modulating immune responses (Unnikrishnan *et al*., 2017, Tran *et al*., 2021, Bowman & Palmer, 2021). While T7SSb machinery varies in sequence and genomic arrangement across species, common components are cytoplasmic protein EsaB and membrane proteins EsaA, EssA, EssB, and EssC, an ATPase that powers secretion of substrates across the bacterial membrane (Bowman & Palmer, 2021). Another distinguished feature of T7SSb is the presence of canonical T7SS substrate WXG100 protein EsxA (named for its 100 amino acid sequence and central Trp-X-Gly [WXG] motif), other small ɑ-helical proteins, and T7SS-associated toxins (Warne *et al*., 2016, Bowran & Palmer, 2021). These toxins, which sometimes contain an N-terminal LXG motif, encode unique C-terminal toxin domains and therefore have biochemically diverse functions. Some T7SS toxins function in interbacterial competition and are frequently co-transcribed with chaperones that facilitate their secretion and immunity factors that prevent self-toxicity. These functions have been described in *Staphylococcus aureus, Streptococcus intermedius, Bacillus subtilis, Enterococcus faecalis*, and recently *Streptococcus gallolyticus* (Cao *et al*., 2016, Whitney *et al*., 2017, Klein *et al*., 2018, Ulhuq *et al*., 2020, Kobayashi, 2021, Chatterjee *et al*., 2021, Teh *et al*., 2022, Klein *et al*., 2022). Interestingly, in some cases, these toxins also promote virulence and modulate immune responses within the host (Dai *et al*., 2017, Ohr *et al*., 2017, Ulhuq *et al*., 2020).

Despite these common traits, the T7SSb encodes largely unique effectors across species. Extensive intra-species T7SS diversity has also been characterized in *S. aureus, Listeria monocytogenes*, and *Staphylococcus lugdunensis* T7SSb based on EssC C-terminus variants and downstream effectors (Warne *et al*., 2016, Lebeurre *et al*., 2019, Bowran & Palmer, 2021). These EssC variants are thought to determine specificity for substrate recognition and secretion. In *S. aureus*, although all four EssC variants are capable of secreting common T7SS substrate EsxA, the EssC2, EssC3 and EssC4 variants are incapable of secreting non-cognate substrate EsxC (which is encoded for downstream of *essC1* only) (Jager *et al*., 2018). We recently showed intra-species diversity in T7SS downstream effectors in group B *Streptococcus* (GBS) (Spencer *et al*., 2021b), and hypothesized that variant-specific T7SSb effectors may promote differing bacterial interactions with the host or other microbes (Spencer & Doran, 2022).

As an opportunistic pathogen, GBS (or *Streptococcus agalactiae*) asymptomatically resides in the gastrointestinal and/or female genital tract of 25-30% of healthy adults (Wilkinson, 1978, Regan *et al*., 1991) but can cause severe infections in some individuals, such as pregnant people and newborns, the elderly, and patients living with cancer or diabetes (Nandyal, 2008, Pimentel *et al*., 2016, Russell *et al*., 2017, Patras & Nizet, 2018, van Kassel *et al*., 2019, Navarro-Torne *et al*., 2021). Within the female genital tract, GBS coexists and/or competes with the vaginal microbiota and, therefore, has evolved mechanisms to survive these encounters while also avoiding immune clearance (Okumura & Nizet, 2014, Vrbanac *et al*., 2018, Coleman *et al*., 2021). We showed previously that GBS T7SS subtype I plays a role in virulence, cytotoxicity, and pore formation via the secreted effector EsxA (Spencer *et al*., 2021b), and that four different GBS T7SS subtypes can be delineated based on differing number of copies of *esxA*, a unique EssC ATPase C-terminus, and a unique repertoire of downstream genes/putative effectors. However, the full heterogeneity of the GBS T7SS operon and the functions of the putative T7SS effectors in GBS-host and GBS-microbe interactions has not yet been investigated.

We hypothesized that GBS T7SS subtypes encode unique downstream effectors that differ from previously studied T7SS substrates in other species and may modulate GBS fitness within the host. Using bioinformatic analyses of the T7SS locus across the four subtypes, we found that the GBS T7SS encodes putative effector proteins with high genetic variability but with similar domains and motifs, suggesting some conserved functions. Functionally, we confirmed that GBS T7SS subtype I and III isolates secrete EsxA *in vitro* and we observed variable impacts of *essC* deficiency across T7SS subtypes *in vivo* using a murine model of female genital tract colonization. This study highlights T7SSb diversity across GBS strains and indicates that the T7SS subtype-specific effector repertoire may differentially modulate host phenotypes.

## 2 RESULTS

### 2.1 Bioinformatic analysis of GBS T7SS subtypes I-IV

In our previous analysis of GBS whole-genome sequences, we identified four GBS T7SS subtypes based on variation in the EssC C-terminus, T7SS substrate EsxA, and several genes encoded downstream of *essC* (Spencer *et al*., 2021b). However, in other species, T7SS loci can encode 10-20 or more genes following *essC* (Bowran & Palmer, 2021, Lebeurre *et al*., 2019, Warne *et al*., 2016). To investigate the full range of GBS T7SS diversity, we extended bioinformatic analysis to the entire T7SS region encoded by 80 T7SSb^+^ GBS genomes deposited in GenBank. Using CJB111, 2603V/R, CNCTC 10/84, and COH1 as example strains for GBS T7SS subtypes I, II, III, and IV respectively, we analyzed each gene within the putative T7SS locus for conserved domains [via InterPro (Blum *et al*., 2021) and Conserved Domain Architecture Retrieval Tool [CDART] analysis (Geer *et al*., 2002)], for predicted protein topology [Protter analysis (Omasits *et al*., 2014)], and for T7SS-associated motifs [(W/F/L)xG and [Y/F]xxxD/E] (**Supp. Table 1**). As discussed in detail below, we found high conservation of GBS T7SS genomic location and machinery between subtypes, but variability in the copy number of locus-associated and orphaned *esxA* genes and heterogeneity in the EssC ATPase and associated putative effector repertoire (**Fig. 1**). We further assessed homology of the T7SS locus to eight other Gram-positive T7SSb-containing species (**Supp. Fig. 1** and **Supp. Table 2**).

**Fig. 1.**
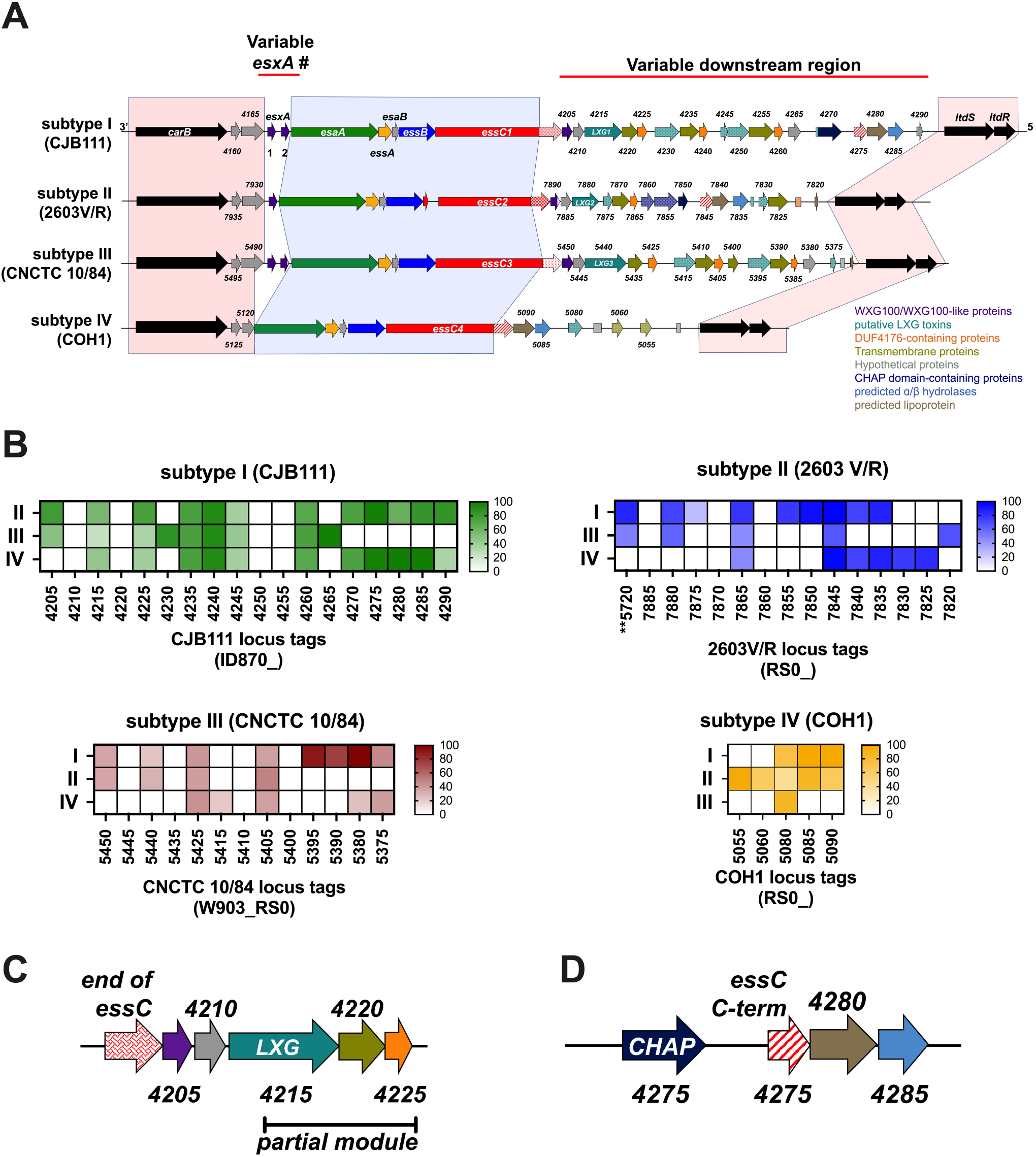
Intra-species diversity of the GBS T7SS. **A)** Example GBS T7SS loci by subtype (roughly to scale). GBS T7SS loci are flanked by carbamoyl phosphate synthesis genes and two-component system genes *ltdS/ltdR* (light red shading). GBS T7SS encode conserved machinery genes (light blue shading) but subtypes vary in copies of *esxA* (purple) and in putative downstream effectors, including WXG100-like proteins (purple), putative LXG toxins (teal), transmembrane proteins (olive green), hypothetical proteins (gray), DUF4176 proteins (orange), CHAP domain-containing proteins (navy blue), ɑβ hydrolases (light blue), and lipoproteins (light brown). Arrows with patterns indicate fragmented genes. Incompletely filled arrows indicate genes encoding prematurely truncated products (e.g., CNCTC 10/84 *esxA2*). **B)** Heatmaps indicating homology of T7SS-associated genes downstream of *essC* across T7SS subtype. Color intensity based on Geneious alignment grade, which considers query coverage and percent identity. Many downstream GBS T7SS effectors have little homology across subtypes or are entirely T7SS subtype specific. **C)** Commonly occurring “LXG modules” are encoded for downstream of *essC* in subtypes I-III, which include putative chaperones (WXG100-like proteins, purple), a putative LXG toxin (teal), a transmembrane protein (olive green), and a DUF4176 protein (orange). Partial LXG modules were also identified, which contain only a fragmented LXG gene (patterned teal) and the subsequent transmembrane and DUF4176 protein encoding genes. **D)** Another commonly found module within T7SS loci consists of a gene fragment encoding the C-terminal end of the COH1 *essC* (red/white stripe), followed by ɑβ hydrolase (light blue) and lipoprotein (light brown) encoding genes. This module is often preceded by an amidase encoding gene (navy blue).

#### WXG100 proteins and putative GBS T7SS machinery encoding genes

Similar to several other Gram-positive bacteria, most GBS T7SS loci contain putative core component genes (*esxA, esaA, essA, esaB, essB*, and *essC*) followed by putative effector and chaperone-encoding genes (Warne *et al*., 2016, Cao *et al*., 2016, Lai *et al*., 2017, Taylor *et al*., 2021, Chatterjee *et al*., 2021). In GBS subtypes I-III, the T7SS locus begins with gene(s) encoding the canonical substrate, WXG100 protein EsxA, which has been hypothesized to also function as a core machinery component of the T7SSb (Sundaramoorthy *et al*., 2008, Kneuper *et al*., 2014). GBS subtype I and III isolates encode two copies of *esxA* (Spencer *et al*., 2021b), while subtype II encodes one copy, and subtype IV strains do not encode a locus-associated *esxA* (**Fig. 1**). The EsxA sequence is highly conserved across GBS T7SS subtypes (≥94% identity) and both EsxA1 and EsxA2 contain the canonical central WXG motif (Spencer *et al*., 2021b) (**Supp. Table 2**).

Downstream of *esxA* (or an WxcM-domain protein encoding gene in subtype IV), putative GBS T7SS machinery genes include *esaA, essA, esaB, essB*, and ATPase-encoding *essC*. These first four core components are highly conserved across GBS T7SS subtypes I-IV, with 97-100% identity at the protein level (**Supp. Table 2**) and appear in the same genomic arrangement as *S. aureus* and *L. monocytogenes* T7SS loci (Kneuper *et al*., 2014, Bowran & Palmer, 2021). Despite this similarity in arrangement, T7SS core machinery sequences exhibit extremely low homology across a panel of Gram-positive species. For example, GBS EsaA is only ∼15-20% identical to EsaA homologs in *Bacillus, Enterococcus, Staphylococcus*, and *Listeria spp*., and only 30-40% identical to streptococcal EsaA homologs in *S. gallolyticus, S. intermedius*, and *S. suis*. Similarly, GBS EssA, EsaB, and EssB proteins exhibited only 11-34% identity to homologs in other genera and 21-54% identity to homologs in other streptococci. The EssC ATPase exhibited the highest inter-species identity, but the lowest intra-species protein sequence identity of the T7SS core proteins. For example, GBS EssC variants were 34-48% identical to non-streptococcal EssC proteins, and 56-80% identical to other streptococcal EssC proteins (**Supp. Table 2**). Within GBS, EssC variants shared 89-98% identity, with sequence variation primarily restricted to the EssC C-terminal 225 amino acids (Spencer *et al*., 2021b). As these variants are associated with unique effector repertoires, it is likely that a given GBS EssC may only export substrates encoded by their cognate subtype, as has been demonstrated in *S. aureus* (Jager *et al*., 2018).

#### Putative T7SS effectors, chaperones, and immunity factors

As many coding sequences within the hypervariable T7SS effector region are annotated as hypothetical, we compared these proteins across GBS subtypes using InterPRO, NCBI CDART, and Protter transmembrane analysis (**Supp. Table 1**). Subtypes I-III encode SACOL2603 T7SS effector proteins (DUF3130-containing or TIGR04197 family proteins), putative LXG toxins, DUF4176-containing proteins, and transmembrane proteins. In addition, amidase domain-containing proteins, C-terminal fragments of EssC, putative toxin fragments, and lipoproteins are also found in some, but not all, GBS T7SS loci (**Fig. 1, Supp. Table 1**). Although similar domains and motifs are detected in different GBS T7SS loci, many proteins differ significantly in sequence homology across subtypes I-III (**Fig. 1B**), suggesting they are distinct proteins with possibly biochemically diverse activities. Interestingly, subtype IV strains do not encode locus associated *esxA* or many common T7SS effectors (SACOL2603, DUF4176, etc.) and encode the shortest GBS T7SS locus, with just 5 hypothetical genes downstream of *essC* that share the highest homology with the subtype II locus (**Fig. 1B**).

Within GBS subtypes I-III variable regions, a commonly occurring module consists of two adjacent WXG100-like proteins (the first containing DUF3130 and the second predicted to contain coil domains), a putative LXG toxin, and one or two hypothetical protein(s), followed by a DUF4176-domain containing protein – hereafter termed an “LXG module” (**Fig. 1C**). A similar gene cluster was recently described in *S. intermedius* where the DUF3130 gene and the adjacent gene encode accessory proteins/chaperones required for secretion of the LXG toxins TelC and TelD (Klein *et al*., 2022). Consequently, these chaperones were named LXG-associated α-helical proteins (Lap). Genes encoding DUF4176 proteins were also observed near this LXG module in *S. intermedius*, although a role for these proteins in T7SS has yet to be shown in any species. A second commonly occurring module in GBS loci consists of a fragment of the subtype IV *essC*, followed by lipoprotein and ɑβ hydrolase encoding genes (**Fig. 1D**).

The first two GBS WXG100-like proteins encoded within the LXG module, hereafter termed putative Lap1 and Lap2, are small and ɑ-helical but lack the canonical central WXG motif; however, Lap1 does display a central FXG motif and a FxxxD/E motif in many GBS strains (**Supp. Fig. 2**). GBS Lap1 proteins exhibit some homology across subtypes I - III (33-75% identity; **Supp. Fig. 2A**) but generally low homology to their Gram-positive counterparts, with ∼40% identity to *S. gallolyticus, S. intermedius*, and *S. suis* (**Supp. Fig. 1**). While neither CDART nor InterPro analyses indicate any putative function for Lap2, these GBS proteins are predicted to have coil domains and to be ɑ-helical in nature (**Fig. 2A, Supp. Table 1**), similar to those described in *S. intermedius* (Klein *et al*., 2022). GBS Lap2 proteins are not well conserved across subtypes I-III, with ≤41% identity between any two (**Supp. Fig. 2D-F**). This indicates that specific Lap/LXG protein pairs may also be required for GBS LXG protein secretion. Although it is currently unknown whether GBS Laps associate with the putative downstream LXG toxin as chaperones, using Alpha Fold modeling and predictions, putative Lap1 and Lap2 across GBS T7SS subtypes seem likely to interact with their cognate LXG protein (**Fig. 2A**) as their counterparts do in *S. intermedius*.

**Fig. 2.**
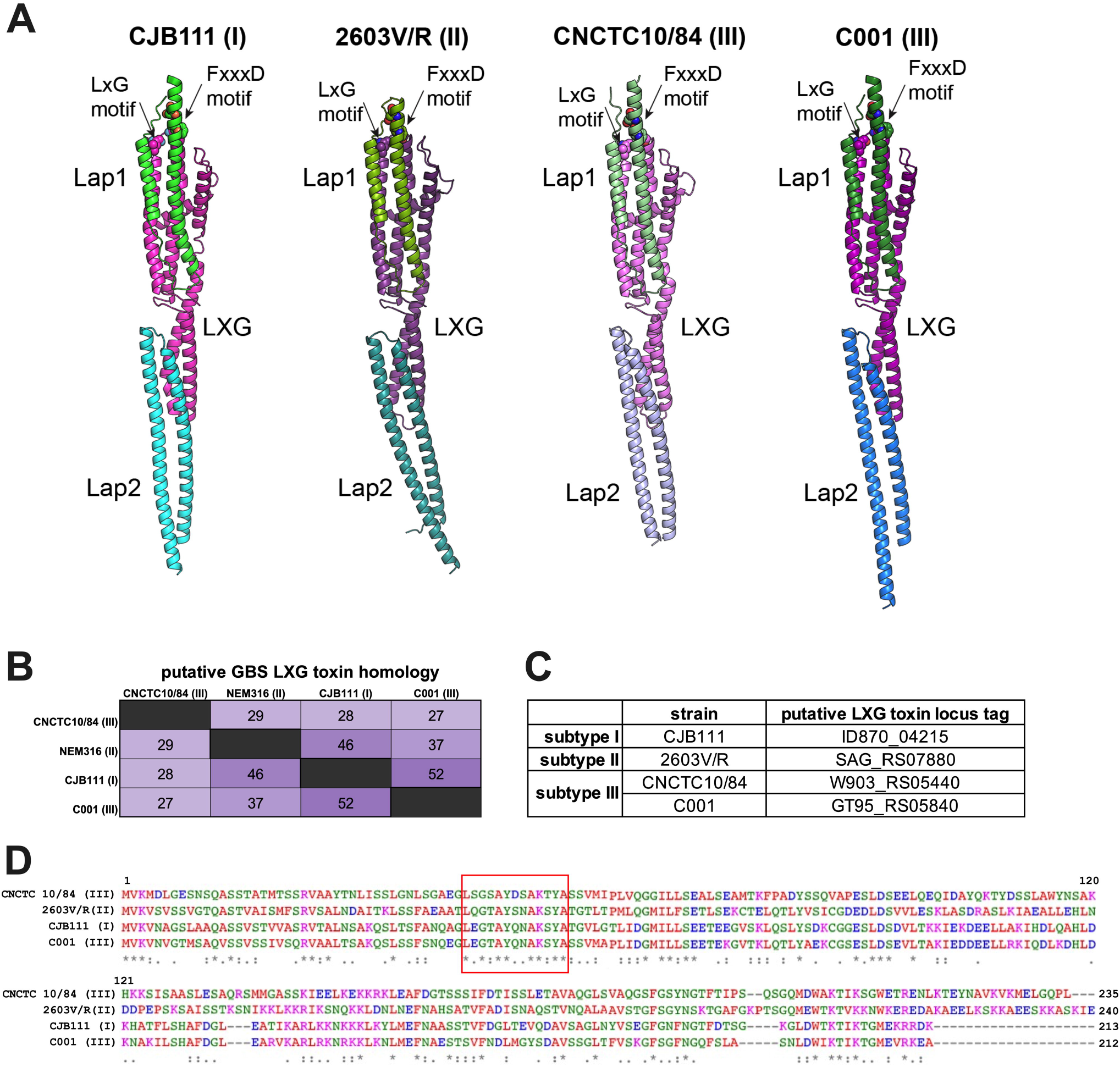
Subtype-specific LXG toxins are encoded by GBS T7SS. **A)** Predicted models of complexes between specific Lap1 (green) and Lap2 (blue/purple) pairs with the LXG domain (pink) of their cognate LXG protein. The putative effector domains of LXG proteins and the predicted disordered regions of Lap proteins are omitted for clarity. The conserved residues of LxG and FxxxD motifs are shown as spheres. The modelling confidence metrics are presented in **Supp. Fig. 3. B-D)** Percent identity matrix of full-length LXG protein sequences and Clustal Omega alignment of the LXG domain from CJB111 (subtype I), 2603V/R (subtype II), CNCTC 10/84 (subtype III), and C001 (subtype III). In the above matrix, the purple shading corresponds to the level of identity between two strains (on a spectrum of 0 to 100% identity), with darker shading indicative of higher percent identity. The longer motif found in all GBS LXG proteins is denoted by a red box in **D**.

Similar to the Lap-encoding genes, GBS putative LXG toxins exhibit minimal identity across T7SS subtypes, with homology concentrating towards the N-terminal LXG domain and sequences diverging in the C-terminal domains (**Fig. 2B-D**) indicating potential differing biochemical properties. We identified four putative full-length LXG toxins encoded within GBS T7SS loci: one in subtype I (CJB111, ID870_4215), one in subtype II (2603V/R, SAG_RS07880), and two within subtype III strains (CNCTC 10/84, W903_RS05440 and C001, GT95_RS05840) (**Fig. 2B-D; Supp. Fig. 4A**). Upon intra-subtype comparisons, subtype I and II LXG proteins were highly conserved across strains. In 44 of the 46 subtype I strains, the LXG protein exhibited 99.8-100% identity, with just one amino acid substitution (A157E) in the N-terminal LXG domain in some strains. Similarly, the subtype II LXG protein demonstrated 99-100% identity across isolates, with just two of the 15 strains encoding a nonsense mutation at amino acid 218 (G218*) resulting in premature truncation after the LXG domain. As subtype III contains only five completely sequenced strains, further assessment of the two full-length LXG proteins was not possible. GBS T7SS LXG toxins are unique from those encoded by other Gram-positive organisms and any limited homology is restricted to the N-terminal LXG domain within the first ∼200 amino acids of the protein. An exception to this was ∼40% homology observed between the CJB111 LXG C-terminus (ID870_4215) and that of *S. intermedius* TelD (**Supp. Fig. 1**). As CDART/InterPro failed to identify domains within the C-terminus of these proteins (**Supp. Table 1**), the functions of GBS LXG proteins remain unknown and must be determined experimentally. Finally, we observed that the four full-length GBS LXG toxins commonly encode a longer LXG motif: LxGxAYxxAKxYA (**Fig. 2D**). This full motif was conserved across other streptococcal LXG proteins from *S. gallolyticus* (TX20005 JGX27_RS02965) and *S. suis* (WUSS351 E8M06_RS09920). Other Gram-positive LXG proteins contain a similar but slightly abbreviated/modified version of this motif (LxGxAYxxA[K/R]), including those encoded by *S. aureus* (TspA), *S. intermedius* (TelC and TelD), *S. lugdunensis* (HKU09-01 SLGD_RS02660), *S. suis* (WUSS351 E8M06_RS09940 and RS09970), and *S. gallolyticus* (JGX27_RS04665, RS08265, RS03950, and RS11360). This longer and more specific conserved motif may facilitate easier identification of streptococcal LXG toxins in the future.

In several species, T7SS toxins are produced in tandem with an immunity factor to prevent self-toxicity. Downstream of the putative LXG toxins, GBS subtypes I and III encode one hypothetical gene followed by a DUF4176 gene and GBS subtype II encodes two hypothetical genes (a LXG protein fragment [7875] and a predicted transmembrane protein encoding gene [7870]) followed by a DUF4176 gene (**Fig. 1**). We hypothesize that one of these downstream genes may function as an immunity factor for each subtype’s unique LXG protein. In support of this, these hypothetical proteins are highly subtype specific and have low homology to their counterparts across GBS T7SS subtypes (**Supp. Fig. 4B**) or to any proteins found in other Gram-positive T7SSb (CJB111 ID870_4220, **Supp. Fig. 1**). Further, no functional domains were identified in these proteins by cDART or InterPro but, across subtypes I-III, all were predicted to contain two to four transmembrane domains (**Supp. Table 1**, see olive green arrows following teal LXG genes in **Fig. 1**).

DUF4176 proteins are also commonly encoded in the vicinity of T7SSb LXG proteins but their roles in T7SSb are unclear. GBS T7SS subtype I, II, and III strains encode two to three loci associated DUF4176 genes, and most encode an orphaned DUF4176 elsewhere in the genome (occasionally fragmented or annotated as pseudogenes) (**Supp. Fig. 4C**). Of the genes downstream of GBS *essC*, DUF4176 exhibited the most overall homology across subtypes (**Fig. 1B**) as well as to proteins in other T7SSb+ Gram-positive bacteria (30-60% identity across several species; **Supp. Fig. 1**). Further, all GBS DUF4176 proteins encode a central “FXG” motif (**Supp. Fig. 4D**). Interestingly, the orphaned DUF4176 proteins encoded by subtype II, III, and IV strains were almost identical (94-100% identity; subtype I’s orphaned DUF4176 is a pseudogene). Of the locus-associated DUF4176 proteins in subtype I strain CJB111, that encoded by ID870_4240 exhibited more homology to the orphan DUF4176 proteins (81-85% identity) compared to its locus-associated DUF4176 proteins (35-48% identity) (**Supp. Fig. 4E**). Using Alpha Fold, we sought to predict if these putative transmembrane proteins or DUF4176 proteins might interact with their cognate GBS LXG proteins. In CJB111, the transmembrane protein encoded for downstream of LXG (by ID870_4220) yielded the highest confidence score (0.8) indicating there may be a stable interaction between this protein pair, but this would need to be confirmed experimentally.

Some T7SS loci encode additional similarly arranged partial LXG modules further downstream, but these additional modules typically encode a fragmented LXG protein (usually retaining only the central linker region or the C-terminal toxic domain (Zhang *et al*., 2012, Chatterjee *et al*., 2021)). We observed fragmented LXG proteins in all subtypes, including subtype I strain CJB111 (ID870_4230, 4245, and 4250) and in subtype III CNCTC 10/84 (W903_RS05415, RS05395), the majority of which were similarly followed by genes encoding for hypothetical transmembrane and DUF4176 proteins (**Fig. 1A**,**C**). Following these partial LXG modules, the furthest downstream regions of T7SS loci are more variable but sometimes contain blocks of T7SS genes that can be found across subtypes (**Fig. 1**), indicating that this region of the putative T7SS locus may be prone to recombination, as recently shown in *S. aureus* (Garrett *et al*., 2022). This region includes genes encoding hypothetical and transmembrane proteins, CHAP domain containing proteins, FtsK domain containing proteins (C-terminal *essC* fragments), lipoproteins, and ɑβ hydrolases (**Supp. Table 1, Fig. 1A**,**D**).

#### Genomic location and genes flanking GBS T7SSb

In all GBS isolates examined, T7SS genes are located between carbamoyl phosphate synthase genes (*carB*; CJB111 ID870_4155) and the LtdRS two component system (Deng *et al*., 2018, Faralla *et al*., 2014) (**Fig. 1**). Two highly conserved genes downstream of *carB* encode for a 107 amino acid hypothetical protein with no predicted function or domains (CJB111 ID870_4160) and a WxcM domain-containing protein predicted to contain ɑβ hydrolase and/or lipase-like domains (CJB111 ID870_4165). While this WxcM protein is prematurely truncated in subtype IV strains, it maintains >97% identity to the subtype I - III homologs within the first 60% of the amino acid sequence. ID870_4160 homologs were not found within other T7SSb containing Gram-positive species, and the GBS WxcM domain containing protein exhibited minimal identity to homologs in *E. faecalis, S. intermedius*, and *Streptococcus suis*, indicating that these genes are fairly specific to GBS.

To confirm the boundaries of the T7SS operon in subtype I strain CJB111, we assessed which genes are co-transcribed within this genomic region by performing RT-PCR using primers spanning each T7SS gene junction. Primers were confirmed using gDNA and non-specific amplification was assessed using no-RT controls. We observed bands for almost every cDNA RT-PCR reaction, indicating that most genes are capable of being co-transcribed; however, *esxA1* consistently appears to be transcribed separately from the rest of the T7SS operon, indicated by a lack of amplicon from cDNA using primers spanning the *esxA1 - esxA2* junction (**Supp. Fig. 5**). Bands were observed between *esxA2 and* machinery genes of the T7SS indicating they are transcribed together, as has been shown in *S. aureus* and *S. gallolyticus* (Kneuper *et al*., 2014, Taylor *et al*., 2021). Further, we observed bands for all gene junctions until the junction spanning the *ltdS* operon for reactions including cDNA as template (but no bands were detected in the no-RT controls), indicating that the genes within the T7SS locus from *esxA2* through the gene upstream of *ltdS* are capable of being co-transcribed. Although, as RT-PCR can detect low levels of transcript, more sensitive methods should be used in the future to assess potential differential regulation within the T7SS locus.

### 2.2 Intra-subtype diversity of the GBS T7SS

While the above bioinformatic analysis assessed diversity of the GBS T7SS between subtypes and between Gram-positive species, we also observed extensive intra-subtype diversity. Within subtype I strains, the region from *carB* through the 5 genes downstream of *essC* (CJB111 ID870_4225) was highly conserved (∼99% nucleotide identity). However, heterogeneity in the form of insertion or loss of gene blocks occurred further downstream (**Fig. 3A**). For example, 12 out of 46 subtype I strains (representative strain CJB111) encode 18 genes downstream of *essC*. In other subtype I strains (12/46, representative strain A909), the T7SS locus is truncated to just 11 genes downstream of *essC* (ID870_4245 through ID870_4285 have been lost). Additional subtype I T7SS loci (10/46 strains) are almost identical to A909 but have a 73 bp deletion after the first DUF4176 encoding gene (which deletes a putative terminator) and a 69 bp deletion in the intergenic region before A909 gene SAK_RS05570. Finally, compared to the A909 T7SS arrangement, a few strains (representative strain Sag153) have undergone further reductive evolution, losing additional genes in the T7SS locus.

**Fig. 3.**
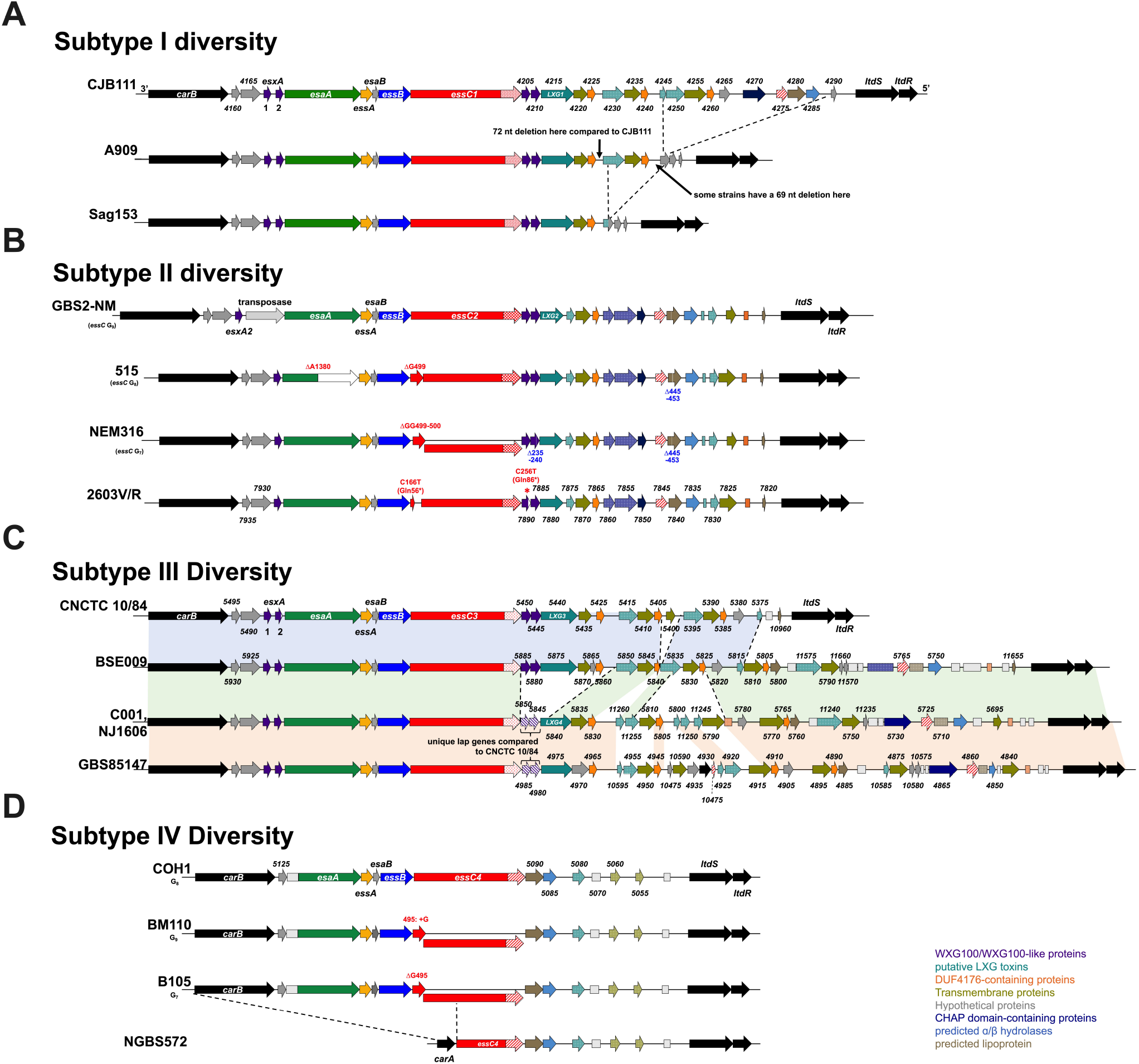
Intra-subtype diversity of the GBS T7SS. GBS T7SS loci representing genomic diversity within **A)** subtype I, **B)** subtype II, **C)** subtype III, and **D)** subtype IV. GBS T7SS encode putative downstream effectors including WXG100-like proteins (purple), putative LXG toxins (teal), transmembrane proteins (olive green), hypothetical proteins (gray), DUF4176 proteins (orange), CHAP domain-containing proteins (navy blue), ɑβ hydrolases (light blue), and lipoproteins (light brown). Arrows with patterns indicate fragmented genes. Incompletely filled arrows indicate genes encoding prematurely truncated products (e.g., CNCTC 10/84 *esxA2*). Subtype I and III strains differ is presence/absence of downstreamgenes **(A, C)**. Subtype II and IV strains differ in mutations within *essC* **(B, D)** and the length of the homopolymeric tract within *essC* is annotated as G_n_.

While most GBS T7SS subtype II strains encode the same set of genes downstream of *essC*, diversity within subtype II is most commonly based on nonsense and frameshift mutations in *essC and esaA* (**Fig. 3B**). Of the 15 closed GBS genomes that encode GBS T7SS subtype II, 13 encode a putatively truncated/multiple-CDS EssC. The majority of these are due to a slip-strand mutation within a homopolymeric G_n_ tract in *essC*, resulting in the loss of either one or two nucleotides, and frameshift and truncation of EssC to 166 or 173 amino acids, respectively. In some subtype II strains, an additional nonsense mutation occurs before this homopolymeric tract resulting in an EssC truncation to 56 amino acids (C166T Gln56*). However, NCBI ORF Finder predicts a second EssC ORF may be possible in these subtype II strains, resulting in a 1291 amino acid protein (of the usual 1469 amino acid protein), which may still allow T7SSb activity. A second common area for mutation within subtype II strains is *esaA* (in 5 of 15 subtype II strains, including strain 515), also potentially due to slippage within a homopolymeric A_n_ tract, resulting in deletion of one nucleotide, frameshift, and truncation of EsaA at 465 amino acids (full length EsaA is 1005 amino acids). Similar to EssC truncation, NCBI ORF Finder predicts a second *esaA* ORF may be encoded; therefore, this mutation may also not necessarily inactivate the T7SS. Despite the T7SS machinery being largely conserved across GBS subtypes (including these homopolymeric tracts), these mutations within T7SS machinery genes rarely occur in GBS subtype I, III, and IV loci.

Subtype III is the least prevalent subtype of GBS T7SS and encompasses immense diversity, with four unique versions of the downstream T7SS region existing in just five isolates (**Fig. 3C**). Similar to subtype I, diversity of subtype III is due to the presence/absence of downstream genes between *essC* and *ltdS*, with some subtype III strains encoding different LXG modules (e.g., strains CNCTC 10/84 and C001 encode different full length LXG proteins). Because these strains are unique in their downstream T7SS gene arrangement, each strain’s downstream repertoire has been independently analyzed in **Supp. Table 1**. Subtype IV strains are also rarer, which impairs intra-subtype comparisons; however, two of 14 subtype IV strains encode similar slip-strand mutations in *essC* as seen in subtype II strains (**Fig. 3D**).

### 2.3 GBS T7SS orphaned WXG100 modules

In addition to locus associated T7SS genes, many GBS strains encode for “orphaned” WXG100 proteins elsewhere in the genome. Comparative genomic analysis of example strains from each T7SS subtype revealed two orphaned WXG100 groups (here termed Orphaned Modules 1 and 2) based on genomic location/neighboring genes and WXG100 protein sequence alignment (**Fig. 4A** and **Supp. Fig. 6**). Module 1 WXG100 orphan proteins are encoded in 64 of the 80 T7SS^+^ GenBank isolate cohort (**Supp. Table 3**), often upstream of a MutR family transcriptional regulator (**Fig. 4A**) and are well conserved, exhibiting 97-100% identity across strains (**Supp. Fig. 6A-B**). Module I WXG100 genes are often encoded either in proximity to a RelE/ParE family toxin protein and plasmid recombinase genes in some strains (as in NEM316) or downstream of a DEAD/DEAH box helicase gene in others (CJB111 and 515). Furthermore, Module 1 WXG100 orphan genes are commonly followed by a cluster of conserved hypothetical genes. In a subset of subtype IV strains, the orphan WXG100 gene is encoded downstream of *rpsI* (sometimes with restriction-modification and abortive infection system genes immediately preceding the WXG100 gene) and this “Module 1b” contains a ∼15kb insertion (encoding for tyrosine recombinases, sigma factors, and *tetM*) following the second hypothetical gene of the cluster downstream of the WXG100 gene.

**Fig. 4.**
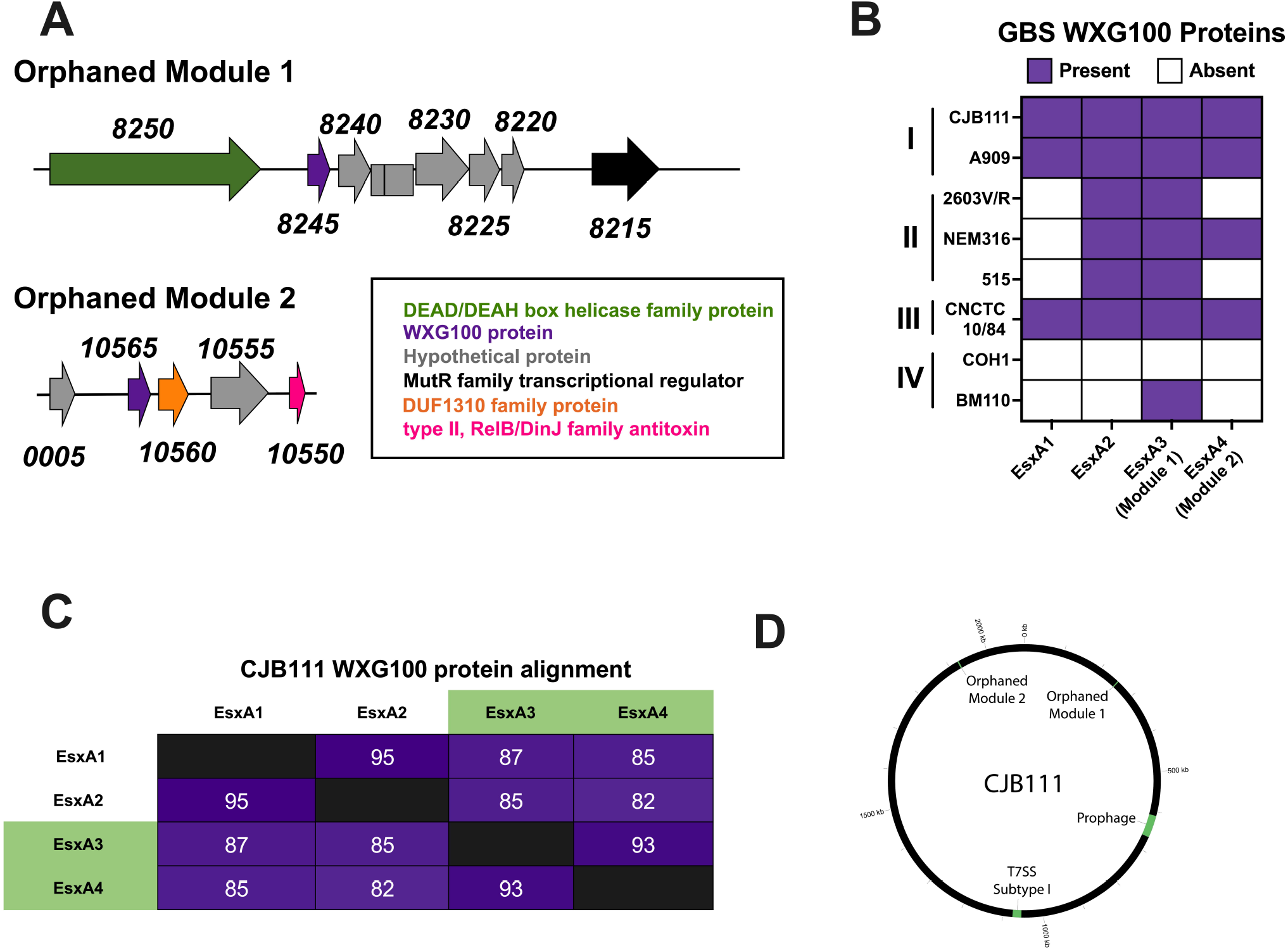
GBS T7SS orphaned modules. **A)** Representative GBS T7SS orphaned modules 1 and 2. The subtype I strain CJB111 Module 1 region includes genes encoding a DEAH/DEAD box helicase (green), WXG100 protein (purple), hypothetical proteins (gray), and a MutR family transcriptional regulator (black). The Module 2 region includes genes encoding hypothetical proteins (gray), a WXG100 protein (purple), DUF1310 protein(s) (orange), and a type II RelB/DinJ family antitoxin (pink). **B)** Heatmap indicating presence (purple) or absence (white) of locus associated EsxA1 and EsxA2 and of GBS T7SS orphaned modules 1 and 2 (containing EsxA3 and EsxA4, respectively) in various GBS clinical isolates. **C)** Percent identity matrix of EsxA1-4 protein sequences from CJB111 (subtype I). In the above matrix, the purple shading corresponds to the level of identity between two strains (on a spectrum of 0 to 100% identity), with darker shading indicative of higher percent identity. **D)** Circos plot showing distinct genomic locations of the T7SS locus, orphaned modules, and prophage in subtype I strain CJB111.

Module 2 WXG100 orphans are less common (encoded in just 45 of the 80 T7SS^+^ GenBank isolates) and are only observed in strains encoding the Module 1 WXG100 orphan (**Fig 4B, Supp. Table 3)**. While the genomic region surrounding Module 2 orphans are more variable, this WXG100 gene (ID870_10565 in subtype I strain CJB111) is consistently encoded upstream of at least one DUF1310 gene and a few genes upstream of a Type II toxin-antitoxin system RelB/DinJ family antitoxin fragment (**Fig. 4A**). Module 2 WXG100 orphans are well-conserved at the protein sequence level (**Supp. Fig. 6C**) and orphaned WXG100 proteins are more similar to each other compared to locus associated EsxA in subtype I strain CJB111 (**Fig. 4C**). However, because orphaned WXG100 proteins are distinct from WXG100 proteins found in other species, we have continued the *esxA* nomenclature and have annotated these genes as *esxA3* (Module 1 orphan) and *esxA4* (Module 2 orphan). Integrase/recombinase genes and toxin-antitoxin system genes are often associated with phage (Harms *et al*., 2018), which are known to mediate horizontal gene transfer and to modulate bacterial fitness and virulence (Borodovich *et al*., 2022, Basler *et al*., 2012, Hobbs & Mattick, 1993); therefore, we investigated whether GBS T7SS genes may be encoded within prophage. Interestingly, neither T7SS loci nor orphaned WXG100 proteins appear within prophage regions (**Fig. 4D**).

### 2.4 EsxA secretion across T7SS subtypes

We next sought to determine GBS T7SS activity across subtypes by measuring EsxA secretion *in vitro* as previously described (Spencer *et al*., 2021b). We observed EssC-dependent secretion in subtype I strain CJB111 and subtype III strain CNCTC 10/84 but could not detect EsxA in the supernatant or cell pellet of subtype II strain 2603V/R (**Fig. 5**). This lack of detection was not due to inability of the anti-EsxA antibody (raised against CJB111 EsxA1) to bind the 95% identical subtype II EsxA2, since it was cross-reactive against subtype II strain NEM316 EsxA in both the supernatant and cell pellet. These results indicate that T7SS may be repressed or that EsxA may be degraded or unstable in the 2603V/R background. Alternatively, T7SS subtype II strains may require different inducing conditions to express and secrete EsxA *in vitro*. Subtype IV strain COH1 served as a negative control in these studies as it does not encode any EsxA homolog.

**Fig. 5.**
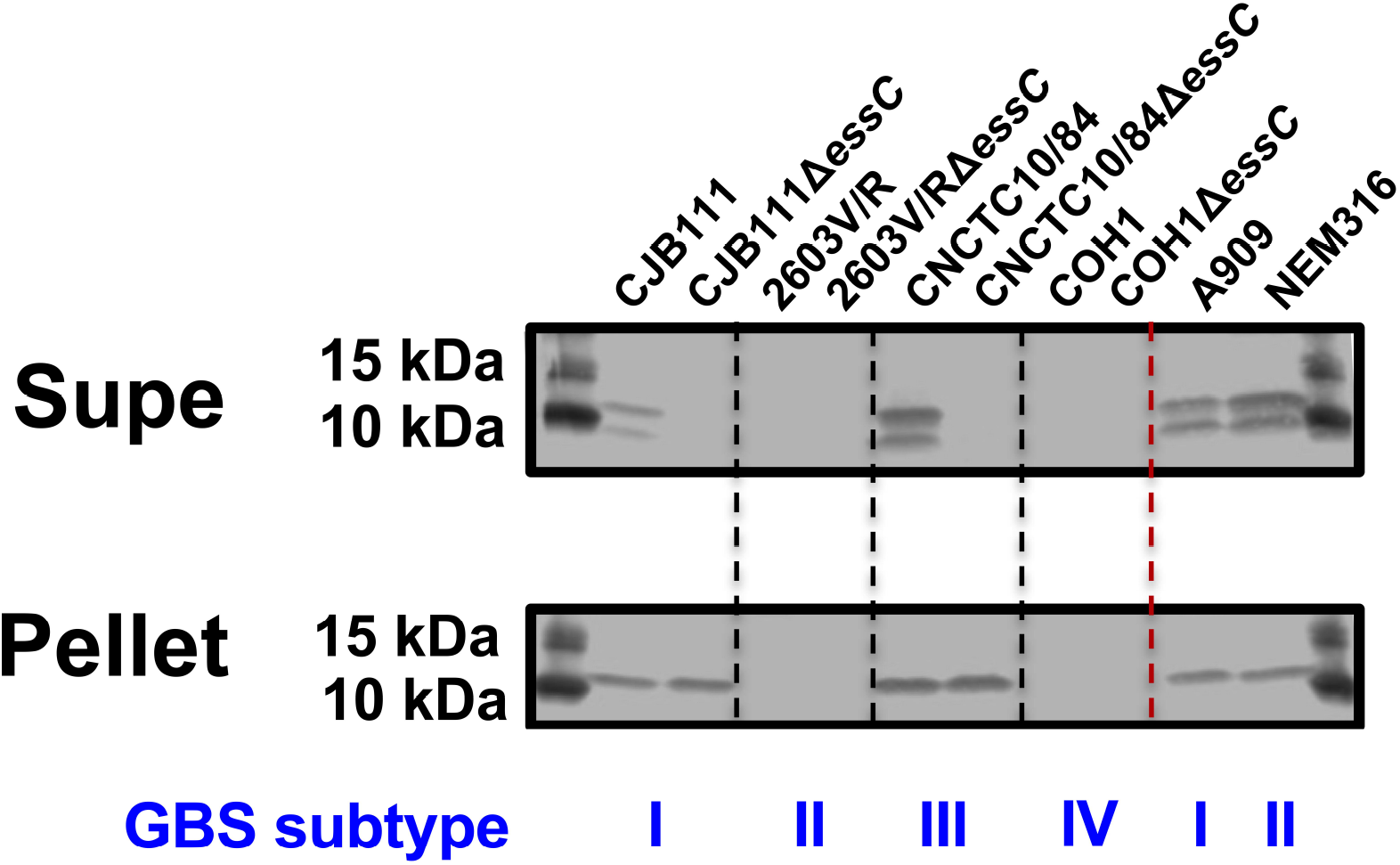
EsxA secretion across GBS T7SS subtypes. Western blot showing EssC-dependent secretion of EsxA from subtype I strain CJB111 and subtype III strain CNCTC 10/84, but not from subtype II strain 2603V/R, *in vitro*. EsxA was also shown to be secreted in subtype I strain A909 and subtype II strain NEM316. Subtype IV strain COH1 does not encode EsxA and served as a negative control. Blot pictured is representative of 3 independent experiments.

### 2.5 Impact of GBS T7SS subtype on vaginal colonization

Secretion systems are often induced by stress, competing bacteria, and/or host pressures. We have previously shown that GBS T7SS promotes systemic infection but hypothesized that T7SS may also play a role during vaginal colonization and ascending infection, niches in which GBS must encounter host immune responses and compete with the native vaginal microbiota. As each GBS T7SS subtype expresses unique putative effectors, we further hypothesized that subtypes may have varying impacts on GBS interaction with the host. As subtypes I and III T7SS are active *in vitro* (as seen by EsxA secretion in **Fig. 5**), we utilized our *essC* deletion mutants in subtype I (CJB111 and A909) and III (CNCTC 10/84) to assess the role of T7SS in vaginal persistence and ascending female genital tract infection in a murine model of colonization. CD1 mice were vaginally inoculated with parental or Δ*essC* mutant strains and GBS persistence over time was assessed by vaginal lavage or swab and tissue burdens were evaluated at the experimental endpoint. Loss of *essC* did not impact subtype I GBS persistence in the vaginal lumen as indicated by percentages of mice vaginally colonized between parental and Δ*essC* colonized groups in CJB111 (**Fig. 6A**) and A909 (**Supp. Fig. 7A**) over time. However, we recovered a marked increase of parental CJB111 from vaginal, cervical, and uterine tissues at the experimental endpoint compared to the CJB111Δ*essC* mutant (**Fig. 6B-D**), indicating that the subtype I T7SS promotes tissue invasion and ascending infection in the female genital tract. Similar trends were observed in the A909 background (**Supp. Fig. 7B-D**). Interestingly, for GBS T7SS subtype III, the CNCTC 10/84Δ*essC* mutant persisted better than the parental CNCTC 10/84 strain in the murine vaginal lumen (**Fig. 6E**). At the experimental endpoint, the CNCTC 10/84Δ*essC* mutant exhibited higher tissue burdens in the vagina, cervix, and uterus compared to parental CNCTC 10/84-colonized mice (**Fig. 6F-H**). Together these data indicate that GBS T7SS subtype I may provide an advantage to GBS in colonization of the female genital tract while T7SS subtype III may be disadvantageous in this environment.

**Fig. 6.**
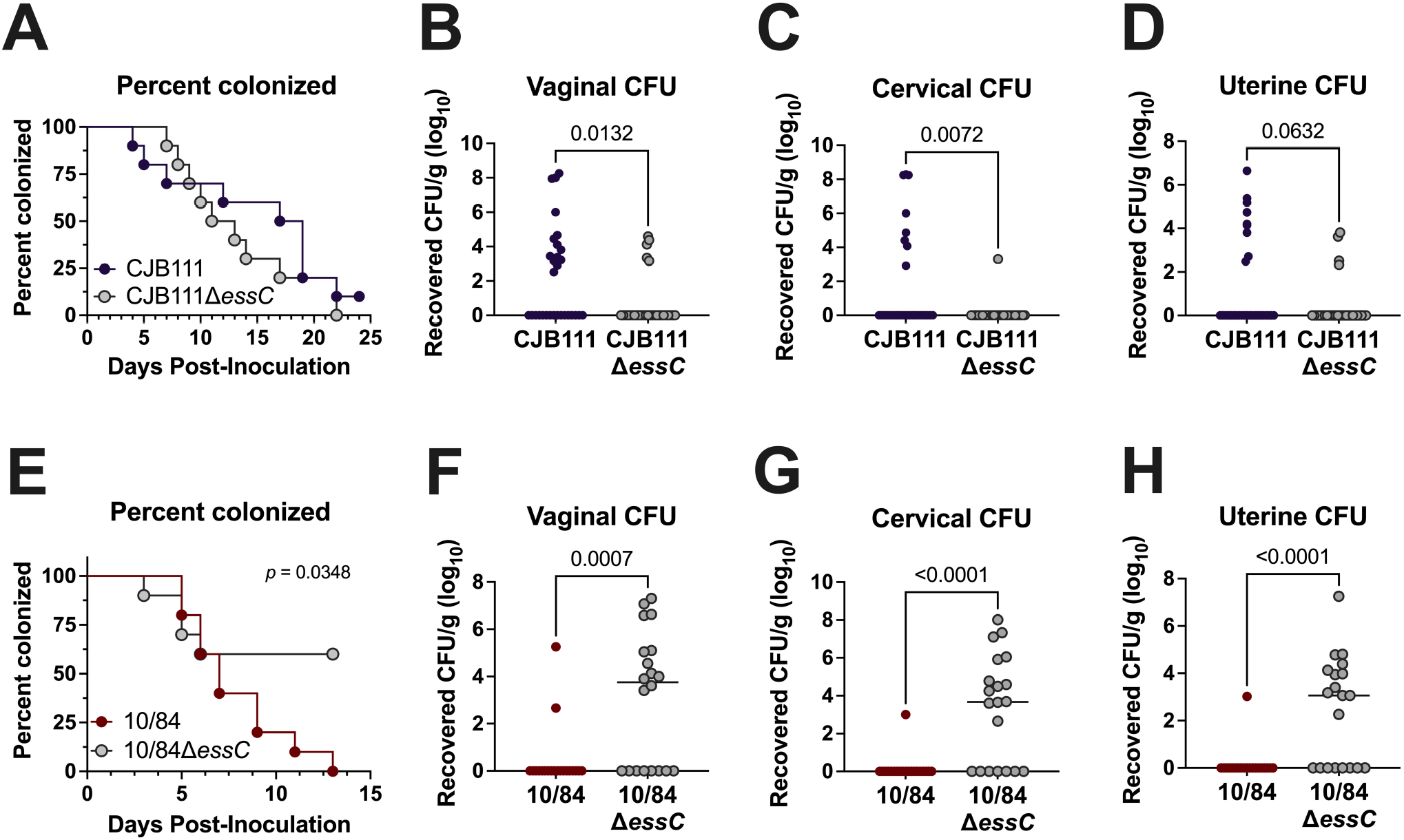
Varying impact of EssC deficiency on GBS vaginal colonization by T7SS subtype I and III strains. **A, E)** Percent colonization curves of 8-week-old CD1 female mice vaginally inoculated with **A)** subtype I strain CJB111 or CJB111Δ*essC* or **E)** subtype III strain CNCTC 10/84 or CNCTC 10/84Δ*essC*. Graphs are representative of three or two independent experiments, respectively (n = 10/group). Statistics reflect the Log rank (Mantel-Cox) test. Recovered CFU counts from the **B, F)** vaginal **C, G)** cervical, and **D, H)** uterine tissue of colonized mice. In panels **B-D** and **F-H**, each dot represents one mouse and all independent experiments’ data are combined in these figures (n = 30/group for CJB111 experiments and n = 20/group for CNCTC 10/84 experiments). Plots show the median and statistics represent the Mann Whitney U test.

### 2.6 Multiplex PCR to T7SS type GBS clinical isolates

Given this striking subtype-specific phenotype, we sought to validate whether the T7SS subtype distribution observed in our GenBank GBS isolates is representative of recent clinical GBS isolates using collections of vaginal isolates from pregnant women and from diabetic foot ulcers that our laboratory has characterized previously (Burcham *et al*., 2019, Keogh *et al*., 2022). To determine GBS T7SS subtypes, we developed a multiplex PCR assay using primers against subtype specific transmembrane encoding genes, the amplicons of which could be distinguished by size (**Fig. 7A**; **Supp. Table 4**). Within the clinical isolate cohorts, most encode T7SS subtype I and subtype II (24/64, 37.5% and 33/64, 51.6%, respectively) (**Fig. 7B-C**). A small percentage of isolates encode subtype IV (7/64, 10.9%); however, we did not identify any subtype III in our clinical isolate cohorts, corroborating its low prevalence in whole genome sequences available on GenBank. We compiled relevant information on these strains as well as those from GenBank including sequence type, serotype, T7SS locus subtype, prophage cluster, and orphan modules encoded to determine if any associations exist between these metrics using Fisher’s exact tests (**Supp. Tables 3 and 5**). Similar to a previous report on GBS T7SS genomic arrangement (Zhou *et al*., 2022) we found that GBS T7SS subtype is associated with sequence-type (*p* < 2.2e-16) and is also associated with GBS serotype (*p* < 2.158E-13). Within our clinical isolate cohorts, trends were particularly observed between T7SS subtype I and serotype Ib, T7SS subtype II and serotypes Ia and II, and T7SS subtype IV with serotype III sequence type 17 (ST-17) (**Supp. Tables 3 and 5**). While we also observed significant associations between prophage cluster, orphaned modules, and T7SS subtype in our vaginal isolate cohort, it is possible these associations may be artifacts of T7SS association with sequence type.

**Fig. 7.**
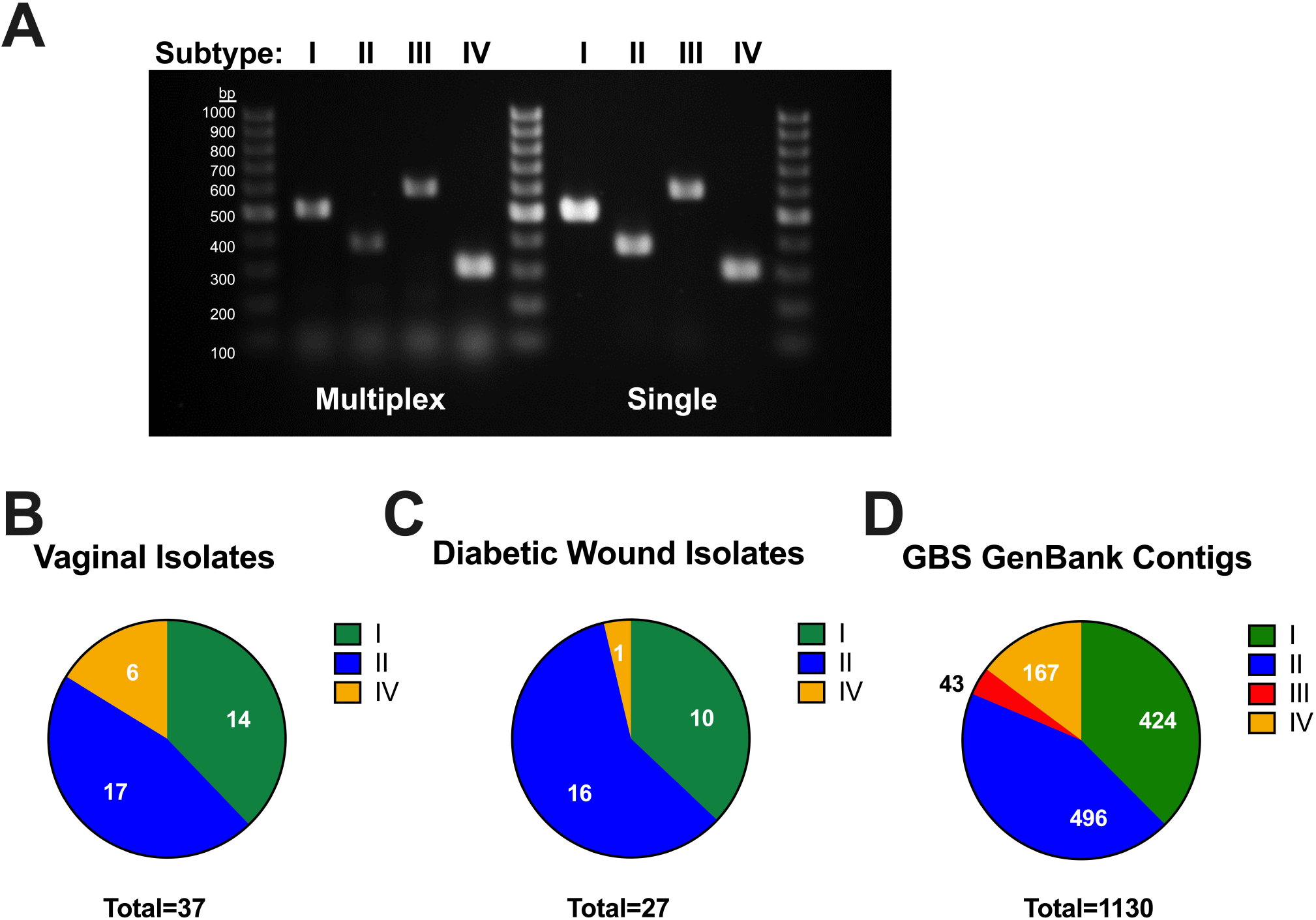
Multiplex PCR to identify T7SS subtype amongst GBS isolates. A multiplex PCR was developed for GBS T7SS typing using primers that amplify subtype specific genes, designed to yield distinct amplicon sizes across T7SS subtypes (see **Supp. Table 4**). **A)** Agarose gel showing subtype specific amplicon products from single-plex and multiplex PCR (of GBS genomic DNA), which can be differentiated by size using a 100 bp DNA ladder. This multiplexed PCR system was utilized in the laboratory or *in silico* using Geneious to T7SS type GBS **B)** vaginal (n =37) and **C)** diabetic wound isolates (n = 27) or **D)** GBS contig sequences available from GenBank (n =1130), respectively.

To achieve a larger dataset and to ensure that our isolate banks are not biased due to collection, we performed *in silico* T7SS subtyping of WGS GBS sequences available in contigs. Of 1342 sequences, 1130 were able to be typed using BLAST of the 225 C-terminal amino acids of EssC and the subtype specific gene used for physical typing of isolates in **Supp. Table 4** and **Fig. 7**. Strains were only considered typable if positive for both a specific subtype’s EssC and downstream subtype-specific gene. Upon screening of these WGS sequence contigs of GBS isolates with typable T7SS, 424 were subtype I (37.5%), 496 were subtype II (43.9%), 43 were subtype III (3.8%), and 167 were subtype IV (14.8%) (**Fig. 7D**). As these sequences are in contigs, T7SS typing was not possible for all isolates (sequences may have gaps within the genomic region containing the T7SS locus or, occasionally, some strains do not encode a T7SS). Sequence analysis of these additional strains further confirmed that subtypes I and II are the most prevalent and that extensive diversity exists within GBS T7SS.

## 3 DISCUSSION

This study highlights the heterogeneity of GBS T7SS subtypes, which could affect GBS interactions in the host. One purpose of this work was to compare GBS T7SS to T7SS loci encoded by other Gram-positive species. Despite having a similar arrangement of T7SS genes, GBS core proteins and putative effectors were largely unique compared to T7SS proteins described in eight other Gram-positive organisms. The secreted substrate/putative core protein EsxA and ATPase EssC were the most conserved T7SS components, consistent with previous observations that these two proteins are general features of T7SS (Pallen, 2002). Despite low sequence homology, GBS T7SS components are commonly found within T7SSb in other species, including WXG-like DUF3130 containing proteins, putative LXG toxins, putative immunity factors and chaperones, CHAP domain containing putative lysins, transmembrane proteins, lipoproteins, fragments of *essC*, and DUF4176 proteins.

Within GBS, T7SS loci reside in a common genomic location, contain homologous machinery, subtype-specific putative effectors/immunity factors/chaperones downstream of *essC*, and, with the exception of subtype IV, encode at least one copy of *esxA* upstream of T7SS machinery. However, we also observed extensive intra-subtype GBS T7SS heterogeneity. Within subtype I specifically, strains encoding the A909 type locus lack approximately eight genes compared to the subtype I CJB111 locus; yet A909 maintains functional EsxA secretion (**Fig. 3A, Fig. 5**). In vaginal colonization experiments, parental CJB111 and A909 exhibited higher tissue invasion compared to Δ*essC* mutants (**Fig. 6B-D, Supp. Fig. 7B-D**). It is unclear whether this eight gene cluster contributes to T7SS activity and future work will investigate its possible contributions to GBS-host interactions. GBS T7SS subtype II loci differ from each other based on truncations of T7SS machinery, particularly within *essC* due to either nonsense or slip strand mutations. However, these putative truncated EssC proteins may retain function as subtype II strain NEM316 is capable of EsxA secretion (**Fig. 5**). It is possible that a second EssC ORF is transcribed and translated, and whether this second CDS works with EssC CDS1 or is functional on its own would be interesting to investigate.

Across species and strains, T7SSb proteins also commonly contain T7-associated motifs, including the [W/F]XG motif (found within EsxA and other WXG100-like proteins), the LXG motif (found within toxins that promote interbacterial competition), and the YxxxD/E motif. The YxxxD/E motif is thought to form one part of a bipartite signal required for homing of a substrate to the machinery for export in mycobacterial T7SSa and Firmicute T7SSb (Champion *et al*., 2006, Daleke *et al*., 2012, Anderson *et al*., 2013, Sysoeva *et al*., 2014). Permutations of these motifs also exist, as a C-terminal FxxxD/E motif was found within GBS Lap1 and “FXG” motifs were identified in GBS Lap1 and DUF4176 proteins. While the presence of a given motif does not necessitate a role in T7SS, we observed that many machinery and putative effector proteins encode YxxxD/E motifs, which may be important for substrate recognition. Interestingly, unlike mycobacterial and staphylococcal EsxA homologs, GBS EsxA1 and EsxA2 do not encode a C-terminal YxxxD/E motif. Previously, Poulsen *et al*. showed that some WXG100 proteins may encode a less specific C-terminal motif, which might direct these substrates to the T7SS machinery: HxxxD/ExxhxxxH (in which the H and h, stand for high and low conservation of hydrophobic resides, respectively (Poulsen *et al*., 2014)). Both GBS EsxA1 and EsxA2 C-terminal sequences include an HxxxD/ExxhxxxH motif, as does the *S. gallolyticus* EsxA (Taylor *et al*., 2021); however, further studies are needed to determine whether this C-terminal motif is required for GBS EsxA secretion.

LXG modules (an LXG gene preceded by two small alpha helical protein genes and followed by one or two hypothetical proteins(s) and a DUF4176 gene) are widespread in T7SSb^+^ Gram-positive species. In *S. aureus*, two WXG-like proteins named EsxC and EsxD can heterodimerize with EsxA and EsxB, respectively, and substrate secretion is interdependent as deletion of these WXG-like genes impacts EsxA/B export (Anderson *et al*., 2013). In *S. intermedius*, WXG-like genes adjacent to *essC* facilitate secretion of a downstream LXG toxin (Klein *et al*., 2022). We have observed that a double mutant in both these WXG-like genes in GBS subtype I did not affect EsxA secretion (data not shown); therefore, we hypothesize that GBS WXG-like proteins may function in LXG protein secretion, similar to *S. intermedius*. Transmembrane protein and DUF4176 protein encoding genes, commonly observed in the vicinity of T7SS loci and toxins, appeared downstream of the LXG genes in GBS subtypes I-III. In *S. aureus*, cognate immunity factor *esaG* (DUF600) is encoded downstream of nuclease EsaD (Cao *et al*., 2016) and in *S. intermedius*, Tel immunity proteins are encoded either adjacent to (in the case of TelB, TelC, TelD) or one gene separated (in the case of TelA) from the LXG toxin (Whitney *et al*., 2017, Klein *et al*., 2022). We therefore expect that one of these downstream GBS genes encodes a subtype specific immunity factor and determining these LXG toxin-immunity factor pairs is the subject of our future work.

In GBS T7SS subtypes I-III, LXG proteins are typically encoded by the third gene downstream of *essC* and are associated with unique upstream genes (putative chaperones) and unique downstream genes (putative immunity factors). These LXG proteins have highly conserved ɑ-helical N-termini within and across GBS subtypes, which are structurally similar to the WXG100 proteins (as originally described by Aravind *et al*., 2011), but have unique, globular C-terminal (and putatively toxic) domains (**Supp. Fig. 4A**). Our bioinformatic analysis failed to identify domains/predicted functions for these C-terminal putatively toxic regions; thus, our future work will determine the putatively toxic activities of these proteins experimentally. Because the toxic activity and chaperone/accessory functions for these proteins have not yet been demonstrated experimentally, we have named these GBS gene products conservatively as LXG-domain containing proteins and the WXG100-like proteins as putative LXG-associated proteins. Many T7SS^+^ Gram positive species also encode orphaned LXG proteins elsewhere in the genome. For example, LXG toxin TspA is secreted by the T7SSb in *S. aureus* but is not encoded in the T7SS locus (Ulhuq *et al*., 2020). Further, *L. monocytogenes, B. subtilis, S. gallolyticus, S. intermedius* and *S. suis* encode multiple full-length LXG toxins, not all of which are associated with the T7SS locus (Bowran & Palmer, 2021, Whitney *et al*., 2017, Kobayashi, 2021, Teh *et al*., 2022, Liang *et al*., 2022). While we did not identify any orphaned full-length GBS LXG proteins based on presence of an LXG motif with the first 100 amino acids of the protein, N-terminal homology to other proteins, or by searching specifically in genomic regions that encode orphaned DUF4176 or WXG100 proteins, it is possible that orphaned C-terminal toxin fragments may exist in GBS strains.

GBS subtype I and II were the most commonly identified T7SS subtypes based on multiplex PCR typing of cohorts of clinical isolates or by *in silico* typing of whole genome sequences and contigs. Subtype IV strains, the next most common subtype, do not encode many common T7SS components, such as locus associated WXG100, DUF3130/SACOL2603, or full-length LXG or DUF4176 proteins. While the genes encoded between the COH1 subtype IV *essC* and *ltdS* are all annotated as hypothetical, interestingly, some of these COH1 genes (the *essC*-lipoprotein-hydrolase module depicted in **Fig. 1D**) also appear in subtype I, II, and III loci; see **Fig 1** and **Fig. 3**), indicating that homologous recombination may occur in T7SS loci as has been reported in other species. While the biological purpose of the GBS T7SS subtype IV is currently unclear, we have observed in subtype IV strain COH1 background that T7SS genes are modulated in certain conditions, such as in a *cas9* deletion mutant (Spencer *et al*., 2019), or during incubation with mucins (Burcham *et al*., 2022b)], thus indicating that the COH1 T7SS may play a role during stress. As most subtype IV strains identified are from the hypervirulent ST-17, serotype III lineage, which is associated with neonatal meningitis, they may have lost some T7SS components due to acquisition of additional virulence factors. Lastly, while subtype I T7SS promoted GBS colonization of genital tract tissues, the T7SS encoded by subtype III strain CNCTC 10/84 appeared to be detrimental for vaginal colonization. We propose that these subtype specific phenotypes are likely due to subtype specific effectors encoded by each strain. Our future work will investigate whether this detrimental role for the CNCTC 10/84 T7SS is due to modulation of the vaginal microbiota or due to modulation of immune responses, resulting in parental CNCTC 10/84 clearance. This disadvantage of the CNCTC 10/84 T7SS in the vaginal tract may account for subtype III T7SS’s rarity across GBS isolates. Indeed, subtype III constitutes just ∼4% of T7SSb^+^ GBS contig sequences (*n* =1130).

As is common for many virulence factors, secretion systems in bacteria often are heavily regulated and may be minimally expressed *in vitro*. We have similarly observed that the GBS T7SS is lowly expressed *in vitro*, requiring concentration of supernatant to detect secreted EsxA (**Fig 5**). Yet, we observe striking GBS T7SS-dependent phenotypes in cell infections or in animal models of infection. Notably, it has been shown that staphylococcal T7SS genes were upregulated in the female genital tract (Deng *et al*., 2019) and TN-seq studies in *E. faecalis* OG1RF revealed that T7SS Tn-mutants were underrepresented in the female genital tract (Alhajjar *et al*., 2020, Burcham *et al*., 2022a), further indicating that this system may be induced in host environments. A common regulatory mechanism for bacterial virulence factors such as adhesins, pili, and capsule is phase variation, which can involve transcriptional slippage resulting in expression of a given factor in some environments and under certain conditions (Phillips *et al*., 2019). For example, transcriptional slippage can occur within genes encoding pneumococcal capsule modifying enzymes, facilitating pneumococcal evasion of vaccine elicited antibodies (van Selm *et al*., 2003, Rajam *et al*., 2007, Spencer *et al*., 2017). We observed many homopolymeric tracts within the GBS T7SS, in line with studies of potential slip-strand regulation in GBS (Janulczyk *et al*., 2010). The most common T7SS slip-strand mutation occurred within *essC* in subtypes II and IV, due to a homopolymeric G_n_ tract. This tract occurs within all subtypes; therefore, it is unclear why these *essC* mutations are not observed in subtype I strains. Because these mutations are primarily found in subtype II, it is possible that subtype specific effectors may induce host pressure, therefore necessitating stringent regulation of this the subtype II locus. Although more work is needed to determine if EssC is encoded as two coding sequences or if the true EssC start site is downstream of the homopolymeric tract, we observed numerous homopolymeric tracts within GBS T7SS loci and hypothesize that these may regulate T7SS gene expression. Finally, numerous single and two-component regulators control T7SS loci across Gram positive species, and this has been most extensively studied in *S. aureus* (Bowman & Palmer, 2021). Interestingly, GBS T7SS gene expression is induced upon deletion of *cas9* (Spencer *et al*., 2019), in the presence of mucins (Burcham *et al*., 2022b), in amniotic fluid (Sitkiewicz *et al*., 2009), and other stress conditions including removal of nutrients, exposure to serum, and oxygen deprivation (Avican *et al*., 2021). Therefore, our investigation of a GBS T7SS regulator is ongoing.

Secretion systems are commonly associated with phage, with a well-established connection between T6SS and bacteriophage machinery in particular. A link between T7SS and bacteriophage infection may also exist as T7SS WXG100 and putative toxins have been identified on *Mycobacterium abscessus* prophage (Dedrick *et al*., 2021) and phage infection induces *E. faecalis* T7SS (Chatterjee *et al*., 2021). Although we do not observe T7SS proteins encoded within GBS prophage, we did observe that some GBS T7SS proteins exhibit homology to phage proteins and that many phage proteins contain T7SS-associated motifs. It is known that encoding of prophage can modulate bacterial resistance to bacteriophage infection and antibiotics (Wendling *et al*., 2021). Recently, integration of a temperate phage into *S. aureus* was shown to increase virulence, not by encoding virulence factors itself, but instead due to upregulation of various bacterial virulence factors including EsxA (Yang *et al*., 2022). This was also recently shown in GBS, in which loss of prophage from CNCTC 10/84 modulated gene expression (Wiafe-Kwakye and Neely, *unpublished observation*). Therefore, future investigation of a link between GBS T7SS and prophage is warranted.

In summary, this study bioinformatically characterizes the diversity encoded within the GBS T7SS and indicates that GBS T7SS contains similar, but unique, individual effectors compared to T7SSb in other Gram-positive species. GBS can be classified into four subtypes, which encode unique effectors and appear to modulate GBS T7SS-dependent interactions within the host, such as during vaginal colonization. This study also identifies orphaned modules containing potential T7SS-associated WXG100 proteins. Taken together, this study suggests a “one size does not fit all” approach to studying T7SS as phenotypes and implications for host and interbacterial phenotypes likely depend on the specific effectors encoded across and even within species. Future studies are warranted to further characterize GBS T7SS effectors and their impact on colonization and disease.

## 4 MATERIALS AND METHODS

### 4.1 Bacterial strains

Example strains from each of the GBS T7SS subtypes were used in this study (subtype I-CJB111; GenBank accession CP063198.2 (01-APR-2021 version) (Spencer *et al*., 2021a), subtype II-2603V/R; GenBank accession NC_004116.1 (Tettelin *et al*., 2002), subtype III-CNCTC 10/84; GenBank accession NZ_CP006910.1 (Nizet *et al*., 1996, Hooven *et al*., 2014, Wilkinson, 1977), subtype IV-COH1; GenBank accession NZ_HG939456.1 (Nizet *et al*., 1996, Da Cunha *et al*., 2014)). Further, previously described cohorts of clinical GBS isolates were utilized for molecular T7SS subtyping and analysis of T7SS activity *in vitro*. Vaginal isolates were obtained from Melody Neely from the Detroit Medical Center as described previously (Burcham *et al*., 2019). Diabetic wound isolates were obtained from Elizabeth Grice (University of Pennsylvania) as well as the CU-Anschutz Medical center (Keogh *et al*., 2022). GBS was grown statically in Todd Hewitt Broth (THB; Research Products International, RPI) at 37°C. All strains used in this study can be found in **Supp. Table 4**.

### 4.2 Bioinformatic analysis of GBS T7SS

All comparative genomics were performed in Geneious Prime 2022.0.2 using genomes of *Streptococcus agalactiae* downloaded from NIH GenBank (as described in Supplementary Table 1 of (Spencer *et al*., 2021b)) as well as using 1342 WGS contigs of *Streptococcus agalactiae* downloaded from NIH GenBank (as of December 2020). Protein BLAST was utilized in Geneious to determine presence/absence or conservation of T7SS-associated proteins across GBS subtypes and across other Gram-positive species. Protein sequences yielding a grade of 30% or less were considered not homologous. Protein alignments were performed using the EMBL-EBI (European Molecular Biology Laboratory-European Bioinformatics Institute) Clustal Omega (v.1.2.4). CDART (Geer *et al*., 2002), and InterPro (Blum *et al*., 2021) were used to identify domains within T7SS locus-associated proteins. Protter was used to characterize protein topology (Omasits *et al*., 2014) and amino acid sequences were scanned manually for T7SS-associated motifs.

### 4.3 Modelling of GBS T7SS proteins

The modelling of LXG proteins and Lap chaperones was performed using original or multimer AlphaFold2 weights (Jumper *et al*., 2021, Evans *et al*., 2022) as implemented in ColabFold (Mirdita *et al*., 2022) or using Robetta (Baek *et al*., 2021). Structural illustrations were generated using PyMol (The PyMOL Molecular Graphics System, Version 2.3.5 Schrödinger, LLC. (Schrodinger, 2015)).

### 4.4 Cloning

A clean *essC* deletion mutant in the CJB111 background was described in (Spencer *et al*., 2021b). Additional deletion mutants were created via allelic exchange in this study in the A909, 2603V/R, CNCTC 10/84, and COH1 backgrounds using the temperature sensitive plasmid pHY304 (Spencer *et al*., 2019) and using a gene encoding spectinomycin resistance (*aad9*) in the knockout construct. Second crossover mutants were screened for erythromycin sensitivity and spectinomycin resistance. Strains created in this study, *essC* locus tags for each strain, and primers used in this study can be found in **Supp. Table 4**.

### 4.5 T7SS operon mapping

Mapping of the T7SS operon in the CJB111 strain background was performed similar to that described previously (Kneuper *et al*., 2014, Taylor *et al*., 2021). Briefly, primers were designed to span gene junctions within the putative T7SS locus and surrounding genomic area. These primers sets were used in PCR reactions with cDNA template (50 ng/ reaction) to determine which adjacent genes were co-transcribed. Genomic DNA (50 ng/ reaction) was used as a positive control for the primer sets and no-reverse transcriptase cDNA was generated and diluted equivalently to control for possible genomic DNA contamination of the cDNA. cDNA and no-RT cDNA were generated as previously described (Spencer *et al*., 2021b). Briefly, GBS strains were grown to mid-log (OD_600_ = 0.4-0.6) and RNA was purified using the MACHEREY-NAGEL NucleoSpin kit (catalog# 740955.250) according to manufacturer instructions with the addition of three bead beating steps (30 sec x 3, with one minute rest on ice between each) following the resuspension of bacterial pellets in RA1 buffer + β-mercaptoethanol. Purified RNA was treated with the Turbo DNase kit (Invitrogen, catalog# AM1907) according to manufacturer instructions. cDNA was synthesized using the SuperScript cDNA synthesis kit (QuantaBio, catalog# 95047-500), per manufacturer instructions. All RT-PCR reactions were performed using Q5 polymerase (New England Biolabs) under the following cycling conditions on a Bio-Rad T100 thermal cycler: 98°C, 2-min hot start; 34 cycles (98°C, 10 seconds; 60 °C, 20 seconds; 72°C, 30-second extension/kb); and 72°C, 10-min extension. Primers used for these experiments can be found in **Supp. Table 4**. PCR amplicons were visualized via gel electrophoresis using 1% agarose gels and GeneRuler 1 kb Plus DNA ladder (Thermo Scientific, SM1331). Three independent replicates were performed.

### 4.6 EsxA secretion assay

Secretion of EsxA during growth in THB was assessed for GBS clinical isolates as described previously (Spencer *et al*., 2021b). Briefly, overnight cultures of GBS isolates were sub-cultured into 5 mL of THB and grown statically for 24 hours at 37°C. Bacteria were pelleted at 3214 x *g* for 10 minutes at 4°C. Supernatants were removed from pellets, filtered (Millex Low Protein Binding Durapore PVDF Membrane 0.22µm filters, catalog #SLGVR33RS), and supplemented with an EDTA-free protease inhibitor cocktail (Millipore-Sigma set III, catalog # 539134; 1:250 dilution). Supernatants were then precipitated overnight with trichloroacetic acid (TCA) at 4°C. Precipitated proteins were centrifuged for 15 minutes at 13K x *g* and resulting pellets were gently washed with acetone. Pellets were then centrifuged again at the same settings. Acetone was removed and pellets were allowed to dry before being resuspended in Tris buffer (50 mM Tris HCl, 10% glycerol, 500 mM NaCl, pH 7). Bacterial pellets were washed once with PBS, frozen overnight, and resuspended in Tris buffer + protease inhibitor. Pellets were then bead beaten (2 x one minute) using 0.1mm zirconia/silica beads (BioSpec). Triton-X-100 was then added to lysates at a final concentration of 1% to solubilize membrane proteins and vortexed to mix.

Supernatant and pellet samples were mixed 1:1 with Laemmli buffer + beta-mercaptoethanol, boiled 10 minutes, and run on SDS-PAGE for Western blotting. Proteins were transferred to membranes via the BioRad Trans-Blot Turbo Transfer System (high molecular weight settings). Membranes were washed three times in TBST and blocked in LI-COR’s Intercept Blocking Buffer (catalog# 927–60001) for one hour at room temperature. Membranes were probed with an anti-EsxA1 rabbit polyclonal antibody (0.5 μg/ml; GenScript) in the above LI-COR blocking buffer overnight at 4°C. Following washes in TBST, membranes were incubated with IRDye 680RD goat anti-rabbit IgG (H + L) secondary antibodies from LI-COR (1:10,000 dilution; 1 hour, room temperature; catalog# 926–68071). Following washes in TBST and water, western blots were imaged using the LI-COR Odyssey.

### 4.7 Murine Model of GBS Vaginal Colonization

GBS vaginal colonization was assessed using a previously described murine model of vaginal persistence and ascending infection (Patras & Doran, 2016). Briefly, 8–10-week-old female CD1 (Charles River) mice were synced with beta-estradiol at day -1 and inoculated with 1 × 10^7^ GBS in PBS on day 0. Post-inoculation, mice were lavaged with PBS or swabbed daily, and the samples were serially diluted and plated for CFU counts to determine bacterial persistence on differential and selective GBS CHROMagar [catalog# SB282(B)]. At experimental end points, mice were euthanized, and female genital tract tissues (vagina, cervix, and uterus) were collected. Tissues were homogenized and samples were serially diluted and plated on CHROMagar for CFU enumeration. Bacterial counts were normalized to the tissue weight. These experiments were approved by the committee on the use and care of animals at the University of Colorado-Anschutz Medical Campus (protocol #00316) and at Baylor College of Medicine (protocol AN-8233).

### 4.8 Molecular T7SS typing of GBS clinical isolates

Molecular typing was performed by multiplex PCR amplification using primers within subtype-specific genes: subtype I, CJB111 ID870_4220; subtype II, 2603V/R SAG_RS07870; subtype III, CNCTC 10/84 W903_RS05410; and subtype IV, COH1 GBSCOH1_RS05060. Primers used for these experiments can be found in **Supp. Table 4**. Multiplex PCR reactions were performed using Q5 polymerase (New England Biolabs) under the following cycling conditions on a Bio-Rad T100 thermal cycler: 98°C, 30 second hot start; 35 cycles (98°C, 10 seconds; 59 °C, 30 seconds; 72°C, 30 second extension/kb); and 72°C, 2-minute extension. PCR amplicons were visualized via gel electrophoresis using 1.4% agarose gels and Gene Ruler 100bp DNA ladder (Thermo Fisher Scientific, SM0243).

### 4.9 Data analysis and statistics

Fisher’s exact test were performed to assess associations between T7SS subtype, T7SS orphaned modules, sequence type, serotype, and prophage cluster. Tables and R script used for Fisher’s exact tests can be found in **Supp. Table 5**. For vaginal colonization experiments, statistical analysis was performed using Prism version 9.4.1 (458) for macOS (GraphPad Software, La Jolla, CA, United States). Significance was defined as p < α, with α = 0.05.

## Acknowledgements

We acknowledge the National Summer Undergraduate Research Program for funding Morgan Apolonio’s virtual research project, which contributed to this work. We also thank Haider Manzer for contributing the Circos plot in **Fig. 4D** and Uday Tak for contributing to the Robetta models in **Supp. Fig. 4A**.

## Supplemental Figure legends

**Supp. Fig. 1. Comparison of GBS T7SS effectors to other Gram-positive organisms**. Heat map comparing CJB111 T7SS protein homology to a panel of eight Gram-positive organisms in which the T7SSb has been studied previously: *S. aureus* (variants 1-4), *S. lugdunensis* (variants 1, 2), *Listeria monocytogenes* (variants 1-7), *E. faecalis* (strain OG1RF), *Bacillus subtilis* (PY79), *S. gallolyticus* (strain TX20005), *S. intermedius* (strains B196 and GC1825), and *S. suis* (strains GZ5065 and WUSS351). See **Supp. Table 2** for individual strain information. Heat map color intensity is based on Geneious alignment grade, which considers query coverage and percent identity. Shading of the locus tag numbers on the x-axis corresponds to the key for gene color in **Fig 1**. Most downstream CJB111 T7SS effectors have little to no homology across species.

**Supp. Fig. 2. Subtype-specific LXG-associated proteins are encoded by GBS T7SS**.

Percent identity matrices and Clustal Omega alignments of **A-C)** LXG-associated protein 1 (Lap1) and **D-F)** LXG-associated protein 2 (Lap2) sequences from CJB111 (subtype I), 2603V/R (subtype II), CNCTC 10/84 (subtype III), and C001 (subtype III). In the above matrices, the purple shading corresponds to the level of identity between two strains (on a spectrum of 0 to 100% identity), with darker shading indicative of higher percent identity. Conserved putative T7SS-associated FXG and FxxxD/E motifs are highlighted by red boxes in **C**.

**Supp. Fig. 3. Confidence scores and predicted aligned error for LXG-Lap Alpha Fold models in Fig. 2A. A)** Predicted LXG-Lap complex models shown in **Fig. 2A** but with color corresponding to per-residue confidence level (predicted local distance difference test [pLDDT] score 1-100; pLDDT > 90 are expected to be modelled to high accuracy). **B)** Predicted aligned error for each LXG-Lap complex, with colors indicating the confidence of domain positions (higher predicted error in red, lower predicted error in blue).

**Supp. Fig. 4. Modeling of full-length LXG proteins and homology of downstream putative immunity factor and DUF4176 genes A)** Robetta predicted structures for full-length putative GBS LXG toxins across subtypes I, II, and III. **B)** Percent identity matrix of putative immunity factors (transmembrane domain containing proteins) encoded for downstream of LXG genes in CJB111 (subtype I), 2603V/R (subtype II), CNCTC 10/84 (subtype III), and C001 (subtype III). **C-E)** Percent identity matrices and Clustal Omega alignments of DUF4176 protein sequences from CJB111 (subtype I), 2603V/R (subtype II), CNCTC 10/84 (subtype III), and COH1 (subtype IV). Green highlighting in **C-E** indicate orphaned DUF4176 proteins. A conserved central FXG motif within GBS DUF4176 proteins is highlighted by a red box in **D**. In all of the above percent identity matrices, the purple shading corresponds to the level of identity between two strains (on a spectrum of 0 to 100% identity), with darker shading indicative of higher percent identity.

**Supp. Fig. 5. Transcriptional landscape of the CJB111 T7SS locus**

RT-PCR was performed to evaluate transcriptional organization of the CJB111 T7SS locus. Using cDNA as template (and no RT-cDNA and genomic DNA as negative and positive controls, respectively), PCRs were performed using primer pairs spanning every gene junction in the putative T7SS locus. Primers used for these experiments can be found in **Supp. Table 4**. Agarose gels shown are representative of three independent experiments.

**Supp. Fig. 6. Homology of orphaned GBS WXG100 proteins**.

**A)** Locus tag tables and percent identity matrices of **B)** Orphaned Module 1 WXG100 protein EsxA3 and **C)** Orphaned Module 2 WXG100 protein EsxA4 sequences across a panel of GBS isolates representing T7SS subtypes I-IV. Green highlighting in **A** indicates orphaned WXG100 proteins. In the above matrices **(B-C)**, the purple shading corresponds to the level of identity between two strains (on a spectrum of 0 to 100% identity), with darker shading indicative of higher percent identity.

**Supp. Fig 7. Impact of EssC deficiency on GBS vaginal colonization by T7SS subtype I strain A909. A)** Percent colonization curve of 8-week-old CD1 female mice vaginally inoculated with subtype I strain A909 or A909Δ*essC*. Statistics reflect the Log rank (Mantel-Cox) test. Recovered CFU counts from the **B)** vaginal **C)** cervical, and **D)** uterine tissue of colonized mice. In panels **B-D**, each dot represents one mouse and two independent experiments’ data are combined in these figures (n = 16/group total). Plots show the median and statistics represent the Mann Whitney U test.

