## Supplemental Figures for "Heterogeneity of the group B streptococcal type VII secretion system and influence on colonization of the female genital tract"

### Supplemental Figure 1

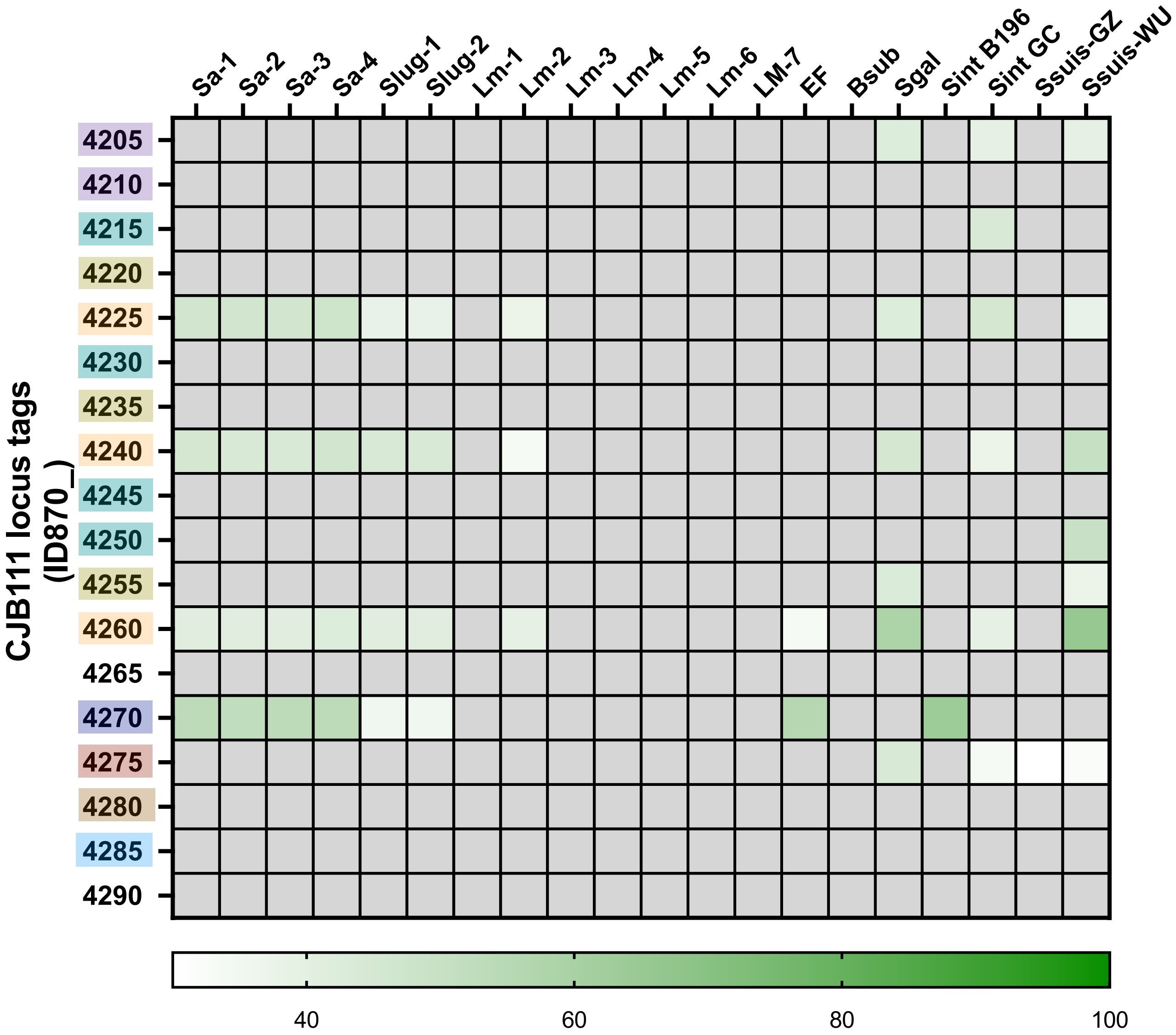


**Shading key:**
WXG100/WXG100-like proteins
putative LXG toxins
DUF4176-containing proteins
Transmembrane proteins
CHAP domain-containing proteins
C-terminal EssC fragment
predicted α/β hydrolases
predicted lipoprotein

### Supplemental Figure 2

## A

**putative LXG associated α-helical protein 1 (TIGR04197 family T7SS effector):**

| | CNCTC10/84 (III) | NEM316 (II) | CJB111 (I) | C001 (III) |
|---|---|---|---|---|
| CNCTC10/84 (III) | | 34 | 41 | 33 |
| NEM316 (II) | 34 | | 62 | 65 |
| CJB111 (I) | 41 | 62 | | 75 |
| C001 (III) | 33 | 65 | 75 | |

## B

| | strain | LAP1/TIGR04197 family type VII secretion effector locus tag |
|---|---|---|
| subtype I | CJB111 | ID870_04205 |
| subtype II | NEM316 | GBS_RS05720 |
| subtype III | CNCTC10/84 | W903_RS05450 |
| | C001 | GT95_RS05850 |

## C

```
CNCTC 10/84 (III) MAKIQSDVSLASSVATGISSAGAEVLSVTKATLDETTTLQGNSTAHEKINLDDATGQQIASSIDTFVGLIHSTASAFVSTDQSLSQMLLNGGEARPTTYKATKPKKKETTLPPAAFNERHTALFGG*
     NEM316 (II) MAKIQSDASVASTCATAIQSGASGITAVGAPTKDSNSDYSGNSDASTYIDGEASSSSQIAEKINEFIALIHSTVNEFEAIDQKISQEIAANTKYT--------MPSNDSSVISN-PDFKPKYGLF*--
     CJB111  (I) MAKIQSDASLASSCATSIQTGANGITSVGKATKDSDSQYSGQTSASLYIDNEASKSGSIAGKLTEFIGLIHSTASEFEAVDQKLSQSISTPKQYT--------TPSGNSTTFQPKPYFN*--------
     C001  (III) MAKIQSNASVASTCATAIQSGASGITAVGKVTKDESSQYSGQTSADLYIDSEASKSVDIAGKLTEFIGLIHSTASEFEAMDSAISQSISVPNQYT--------TNSYTSPSITPKPYFK---------
                ******:.*:**: **.*.:..  : :*   * *. :  .*:: *   *: : :.. .** .:  *:.*****.. * : *. :** :     :             .  .  :      :
```

## D

**putative LXG associated α-helical protein 2**

| | 2603V/R (II) | CNCTC10/84 (III) | CJB111 (I) | C001 (III) |
|---|---|---|---|---|
| 2603V/R (II) | | 15 | 21 | 19 |
| CNCTC10/84 (III) | 15 | | 26 | 24 |
| CJB111 (I) | 21 | 26 | | 41 |
| C001 (III) | 19 | 24 | 41 | |

## E

| | strain | LAP2 locus tag |
|---|---|---|
| subtype I | CJB111 | ID870_04210 |
| subtype II | 2603V/R | GBS_RS07885 |
| subtype III | CNCTC10/84 | W903_RS05445 |
| | C001 | GT95_RS05845 |

## F

```
CNCTC 10/84 (III) MTERTFEDIELDLKLFQIKLDNAENSKRLLQKLKNDVMELQIELLESLKLGDAYLTESE--ELEENNDFILTVNSETLSLEESYDNRINLVSKEIMDYENALDKLYYEKQSLMQKSNERKGG*-------------
      2603V/R(II) MNEKKLSENIEESKKITLEIEKVSDELNQLKLFENDSVDFFDQTQKVF--SDLSLFQKTVRDMSDFEAIRSDYQRLQNLVFIDIDDFKDKLLKKKNQLIQKQDNLYYERQKLQVLPQEKVMKVGEKNGKNGFRRI*
      CJB111  (I) MNDKQLKEVERNEAILRKELERIEDKKIVLKKSYDKTINMQLDIQQSLRDSSQSLSPEE---VMEQEEIMLIFNRQSRIVEDYFQEEMAKLNKQETDAKDTLEGLVQERQKLYVSQSEK-GE*-------------
      C001  (III) MVRMEIEELDNQAATVQKQVEIMEERRLQLEKSYEKAINLQSDIHQSIRNNSLELTQEE---ILDNEEILSVFSRQSRLVEEYYSEEKRQLKEEELKLRDKLDGIIQKRQLLYIENEEK-IRKGELNGKS*-----
                  *    :.:   :   .  :::  .:     *:   :. :::  :  : :  ..  *  .    : : : :    .     :     .:     : ::  .  :  : :  ::* *     .*:
```

### Supplemental Figure 3

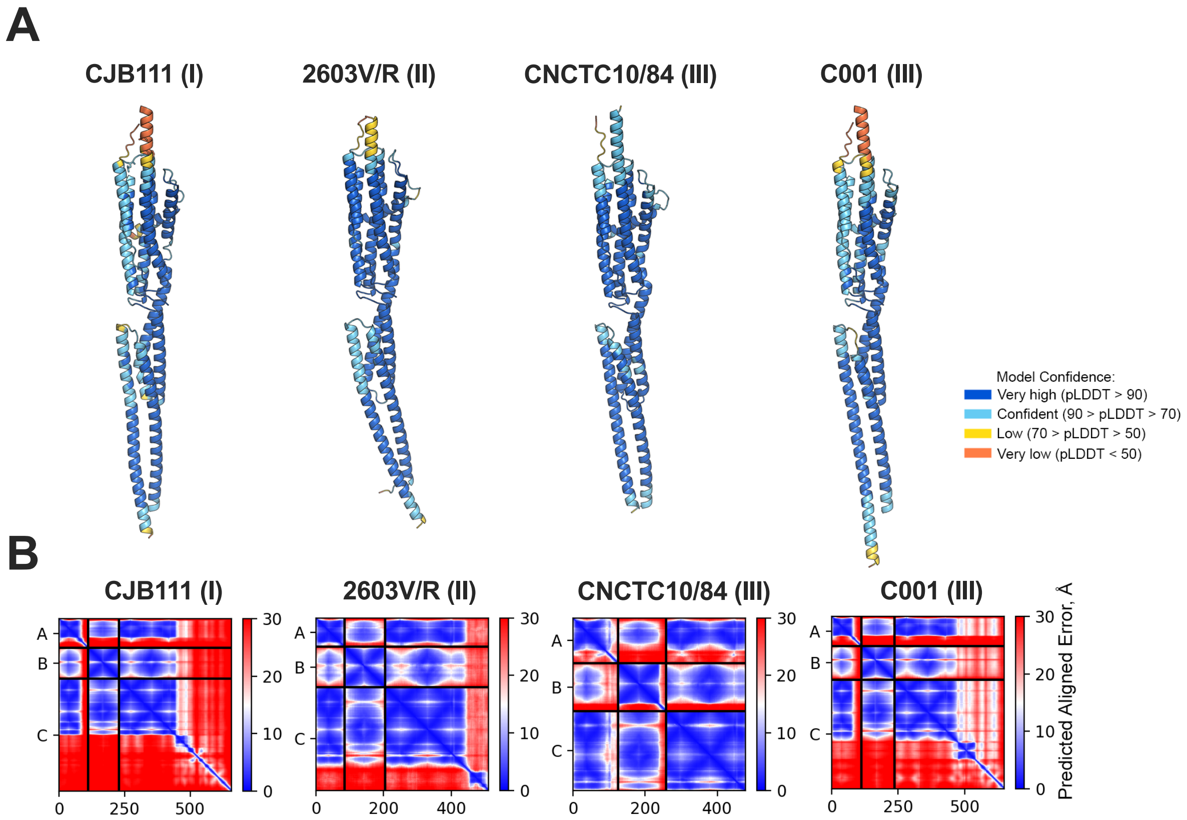

### Supplemental Figure 4

**A**

CJB111 (I) | 2603V/R (II) | CNCTC10/84 (III) | C001 (III)

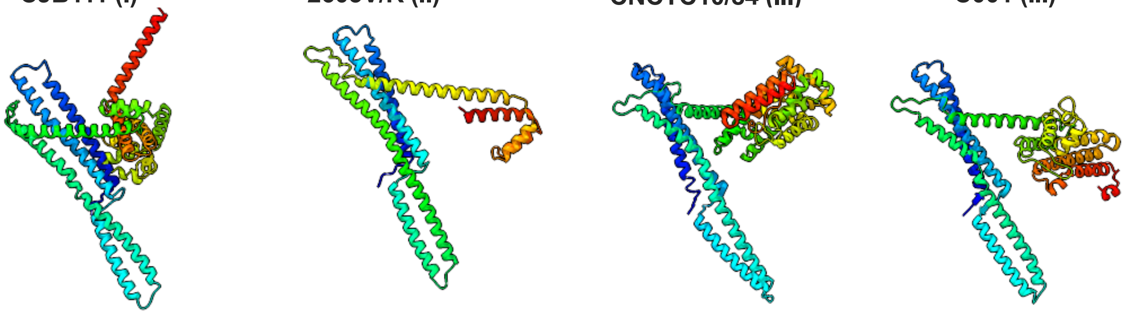

**B**

**Transmembrane proteins downstream of LXG:**

| | CNCTC10/84 (III) | C001 (III) | CJB111 (I) | 2603V/R (II) |
|---|---|---|---|---|
| CNCTC10/84 (III) | | 21 | 20 | 15 |
| C001 (III) | 21 | | 22 | 19 |
| CJB111 (I) | 20 | 22 | | 31 |
| 2603V/R (II) | 15 | 19 | 31 | |

| | strain | downstream genes locus tag |
|---|---|---|
| subtype I | CJB111 | ID870_04220 |
| subtype II | 2603V/R | SAG_RS07870 |
| subtype III | CNCTC10/84 | W903_RS05435 |
| | C001 | GT95_RS05845 |

**C**

#### DUF4176-containing proteins

| subtype I, CJB111 | subtype II, 2603V/R | subtype III, CNCTC 10/84 | subtype IV, COH1 |
|---|---|---|---|
| ID870_RS04225 | SAG_RS07865 | W903_RS05425 | RS08075 |
| ID870_RS04240 | SAG_RS13455 (pseudo) | W903_RS05405 | |
| ID870_RS04260 | SAG_RS11090 | W903_RS05385 | |
| ID870_RS01255 (pseudo) | | W903_RS010490 | |

**D**

```
CNCTC 10/84_DUF2(5405)        --MENKILPLGSIVTLKNGDGSELMIVTRAAIIEED--GVEVYFDYGSVLIPQGMVSPENVYFFNAENVEEVIFKGFENEAEKDFQSNYETMIEQSIYPK---GNVEE---
CNCTC 10/84_DUF1(5425)        --METDNLPIGSVVRLNEGK-IDLIIVSRQPVTVIN--GNKCFFDYAAVSQITGLT-DDKIVFFNNENIEKVIFEGYYGPNEEKVIKAMENWRLNTEIIK---GEIKPSK-
CJB111_DUF1(4225)             ---MEKILPIGSVITLKNGS-LKVMIIARYPLYKYK--NRIGYFDYSGCIFPSGVT-DNQTYFFNNDDIEKVWFTGYIDENELKAQKVFKEQIDKIEYPHFTIEEVNKIEE
2603V/R_DUF1(7865)            ---MDKIHTLGSVIELKNDS-QKVMITSRFPLYDNE--GQLGYFDYSGCIFPISIV-GNETYFFNLEDIDKVLFEGYYDENEEEMQKIFLK--------------------
CJB111_DUF3(4260)             MIKKEELLPLGSIVYLEEGT-VKLMIVGRGVIYDDPEGTGQLFADYMGGLYPAGLQ-TESTLFFQHENIDEVIFEGYRDEEEERFLKIYHEWEKSI-YP------------
CNCTC 10/84_DUF3(5385)        -----------------------MVILNRASIVDQN--GQEVYFDYLGGIFPEGLN-MEQVYYFNEEDIDEVIFEGYHDEEEERFLKIYHEWEKKY-LSLK-VDKITLK--
CJB111_DUF2(4240)             -----MLLPVGSIVYLVDGN-QKLVILNRASIVDQS--GQEVYFDYLGGIFPEGLN-MEQVYYFNEEDIDEVIFEGYHDEEEERVSKLIEKWKQKE-GKNIPKGKTE----
COH1_DUForphan(8075)          -----MLLPVGSVVYLIDGN-QKLVIVNRGAIVEQE--GQEVYFDYLGGIFPEGLN-LEQVYYFNQEDIDEVVFEGYHDEEEERVSRLIEKWKNTE-GKNLPKGKTSVE--
2603V/R_DUForphan(11090)      -----MLLPVGSVVYLIDGN-QKLVIVNRGAIVEQE--GQEVYFDYLGGIFPEGLN-LEQVYYFNQEDIDEVVFEGYHDEEEERVSRLIEKWKNTE-GKNLPKGKTSVE--
CNCTC 10/84_DUForphan(10490)  -------------------------MVNRGAIVEQE--GQEVYFDYLGGIFPEGLN-LEQVYYFNKEDIDEVVFEGYHNEEEERVSRLIEKWKNKE-GKNLPNGKTSVE--
                                                       :  *  :           : ** .     .:   :.  :*: :::::* * *: .   *
```

**E**

| | NCTC_DUF2 (5405) | NCTC_DUF1 (5425) | CJB111_DUF1 (4225) | 2603_DUF1 (7865) | CJB111_DUF3 (4260) | NCTC_DUF3 (5385) | CJB111_DUF2 (4240) | COH1_DUForphan (8075) | 2603_DUForphan (11090) | NCTC_DUForphan (10490) |
|---|---|---|---|---|---|---|---|---|---|---|
| NCTC_DUF2 (5405) | | 33 | 36 | 35 | 40 | 37 | 41 | 40 | 40 | 38 |
| NCTC_DUF1 (5425) | 33 | | 34 | 33 | 35 | 32 | 37 | 36 | 36 | 31 |
| CJB111_DUF1 (4225) | 36 | 34 | | 60 | 35 | 33 | 35 | 33 | 33 | 30 |
| 2603_DUF1 (7865) | 35 | 33 | 60 | | 37 | 48 | 44 | 44 | 44 | 44 |
| CJB111_DUF3 (4260) | 40 | 35 | 35 | 37 | | 56 | 48 | 46 | 46 | 40 |
| NCTC_DUF_3 (5385) | 37 | 32 | 33 | 48 | 56 | | 77 | 64 | 64 | 62 |
| CJB111_DUF2 (4240) | 41 | 37 | 35 | 44 | 48 | 77 | | 85 | 85 | 81 |
| COH1_DUForphan (8075) | 40 | 36 | 33 | 44 | 46 | 64 | 85 | | 100 | 94 |
| 2603_DUForphan (11090) | 40 | 36 | 33 | 44 | 46 | 64 | 85 | 100 | | 94 |
| NCTC_DUForphan (10490) | 38 | 31 | 30 | 44 | 40 | 62 | 81 | 94 | 94 | |

### Supplemental Figure 5

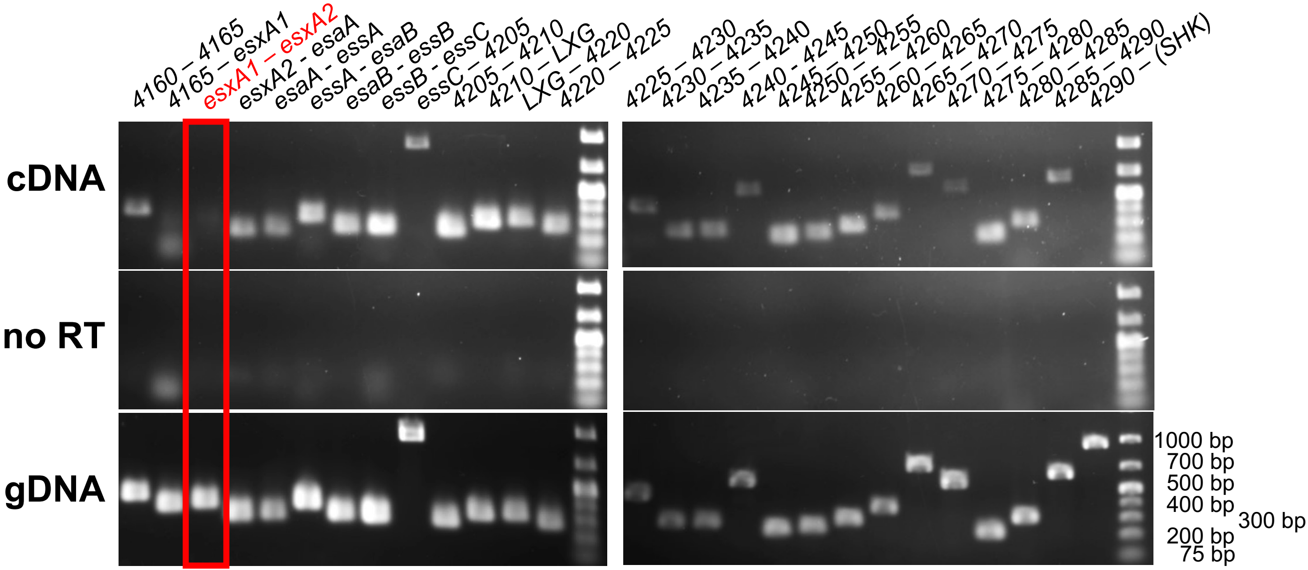

### Supplemental Figure 6

## A

##### GBS WXG100 genes

| | subtype I | | subtype II | | | subtype III | subtype IV | |
|---|---|---|---|---|---|---|---|---|
| | CJB111 (CP063198.2) | A909 (NC_007432.1) | 2603V/R (NC_004116.1) | NEM316 (NC_004368.1) | 515 (NZ_CP051004.1) | CNCTC 10/84 (NZ_CP006910.1) | COH1 (NZ_HG939456.1) | BM110 (NZ_LT714196.1) |
| ***esxA1*** | ID870_04170 | SAK_RS05640 | | | | W903_RS05485 | none | |
| ***esxA2*** | ID870_04175 | SAK_RS05645 | SAG_RS07925 | GBS_RS05755 | GRB95_RS05270 | W903_RS05490 | | |
| ***esxA3*** | ID870_08245 | SAK_RS01430 | SAG_RS03860 | GBS_RS01395 | GRB95_RS01415 | W903_RS01365 | | BQ8897_RS01705 |
| ***esxA4*** | ID870_10565 | SAK_RS09855 | | GBS_RS10285 | | W903_RS10725* | | |

orphaned genes in green

*frameshifted/truncated

## B

##### Module 1 WXG100 (EsxA3) protein alignment

| | CJB111 | A909 | 2603V/R | NEM316 | 515 | CNCTC 10/84 | BM110 |
|---|---|---|---|---|---|---|---|
| **CJB111** | | 100 | 100 | 99 | 99 | 98 | 100 |
| **A909** | 100 | | 100 | 99 | 99 | 98 | 100 |
| **2603V/R** | 100 | 100 | | 99 | 99 | 98 | 100 |
| **NEM316** | 99 | 99 | 99 | | 100 | 97 | 99 |
| **515** | 99 | 99 | 99 | 100 | | 97 | 99 |
| **CNCTC 10/84** | 98 | 98 | 98 | 97 | 97 | | 98 |
| **BM110** | 100 | 100 | 100 | 99 | 99 | 98 | |

## C

##### Module 2 WXG100 (EsxA4) protein alignment

| | CJB111 | A909 | NEM316 | CNCTC 10/84 |
|---|---|---|---|---|
| **CJB111** | | 97 | 99 | 98 |
| **A909** | 97 | | 98 | 95 |
| **NEM316** | 99 | 98 | | 97 |
| **CNCTC 10/84** | 98 | 95 | 97 | |

### Supplemental Figure 7

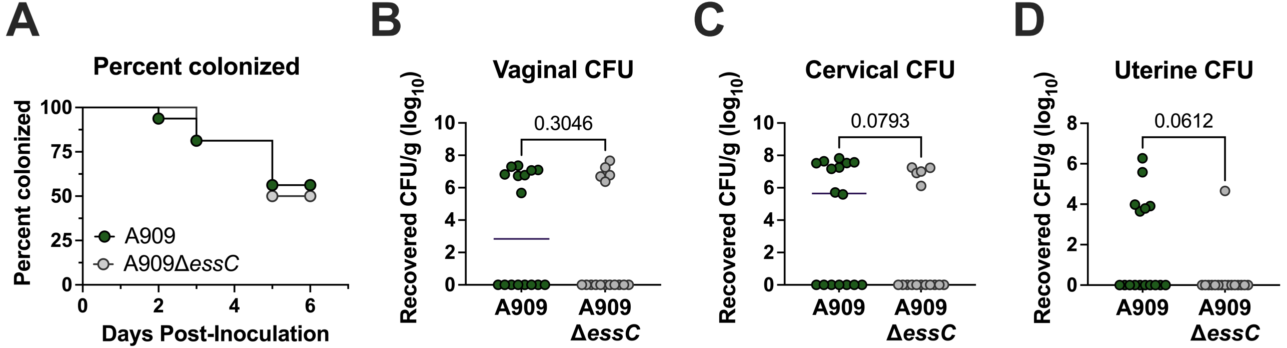
